# Worldwide patterns in mythology echo the human expansion out of Africa

**DOI:** 10.1101/2025.01.24.634692

**Authors:** Hélios Delbrassine, Massimo Mezzavilla, Leonardo Vallini, Yuri Berezkin, Eugenio Bortolini, Jamshid Tehrani, Luca Pagani

## Abstract

Similarities between geographically distant mythological and folkloric traditions have been noted for a long time. With the elaboration of large banks of data describing the presence and absence of narrative motifs around the world, scholars have been able to statistically investigate their potential routes and mechanisms of diffusion. However, despite genetic data allowing for increasingly refined demographic movement inferences, few have integrated it into their models, and none at a global scale. In this work, we capitalise on the augmenting availability of modern and ancient genetic data and on Yuri E. Berezkin’s database of more than 2000 mythological motifs worldwide to investigate the mechanisms involved in generating their present-day distribution at a global scale. The direct combination of both kinds of evidence allows us to explore in more depth the respective influences of population movement and replacement versus cultural diffusion on motif transmission. Our results show that both processes have played important roles in shaping their present-day distribution. By leveraging available ancient DNA (aDNA) and deepening the temporal scale of the detected signals, we reveal that correlations between mythemes and genetic patterns can be traced back to population movements that pre-date the Last Glacial Maximum and go back to at least 38,000 years ago, and possibly even earlier to the human expansion out of Africa some 60,000 years ago. Our work shows the earliest evidence for the transmission of stories and storytelling in human history, and supports the joint use of cultural evolutionary theory and population genetics to illuminate the biocultural processes that shaped our species.

## Introduction

Similarities between geographically distant mythological and folkloric traditions have been noted for a long time^1–37^. Several schools have sought to explain them in various ways, such as psychoanalysis ^18,27^, and structuralism ^21,28^. The identification of basic constituent units to myths and tales and their classification into large databases ^5,17,25,29^ has notably allowed comparative mythologists to investigate their origin through statistical approaches and others developed in the context of evolutionary biology ^16^, such as phylogenetics ^7,8,30–34^. Other approaches have been inspired by population genetics, which are able to capture and quantify cultural admixture between populations more easily than branching phylogenetic structures can. For example, Wright’s F-statistic ^35^ does not require specific units of measurements and can therefore easily be applied to cultural data ^36,37^. Measuring ΦST on folktale distances, Ross et al. (2013)^38^ and Bortolini et al. (2017)^39^ respectively demonstrated significant population structure across ethnolinguistic groups in Europe and in Eurasia.

While these studies demonstrate the rich potential of applying population genetics techniques to folkloristic data, to date there have been few attempts to explicitly integrate the diffusion of traditional narratives with genetic data, despite the opportunities afforded by advances in software and DNA sequencing ^39–43^, with a couple of exceptions. One of these studies ^20^ compared the spatial distribution of American folkloric motifs with that of mitochondrial haplogroups and proposed a Ural-Amerindian mythological complex as ancient as 10-13 Ky old. Focusing instead on Eurasia, Bortlini et al. (2017) ^39^ examined the processes by which folktales had spread, testing how well models of demic and cultural diffusion fit tale distribution patterns. In their work, cultural diffusion was defined as the spread of cultural motifs independently from population movements and replacements (e.g. through word of mouth), while demic diffusion implied the simultaneous turnover of cultural motifs and people (e.g. through range expansions). Overall, their results suggest the predominant effect of language-biased cultural diffusion in populations connected by distances of at least 4,000 km and of demic processes under this range.

In this work, we capitalise on the increased availability of modern and ancient genetic data and on Yuri E. Berezkin’s large database of worldwide mythological and folkloric motifs ^5^, which reports the presence and absence of more than 2000 “mythemes” (defined as concepts, images, or other stable narrative features) in almost 1000 traditions (as of 2018), to study their mechanisms and potential times of diffusion. To do so, we compute mythological distances and correlate them with genetic ones on the one hand (to test for a demic signal) and with geographical ones on the other (to test for cultural dissemination), adapting the methodology developed by Bortolini et al. (2017)^39^. By applying it to various geographical, mythological, and temporal ensembles, we aim to assess the time depth of the processes explaining present-day mytheme distribution worldwide.

## Results

### Admixture between Mythological Traditions

We use ADMIXTURE ^44^ to describe the presence and absence of 2138 mythological motifs in 781 worldwide populations as the combination of nine components (Fig. S2). As one of them is present on all continents (Fig. S3), though in particular in the Southern Hemisphere (Fig. S4), we mask it and rescale the others accordingly. We note that these largely correspond to broad world regions (Fig. 1, Fig S.5), either suggesting a certain time depth allowing for the formation of a cross-continental pattern or, on the contrary, a very recent and independent origin of motifs, with local diffusion being the main driver of the observed pattern.

**Figure 1.**
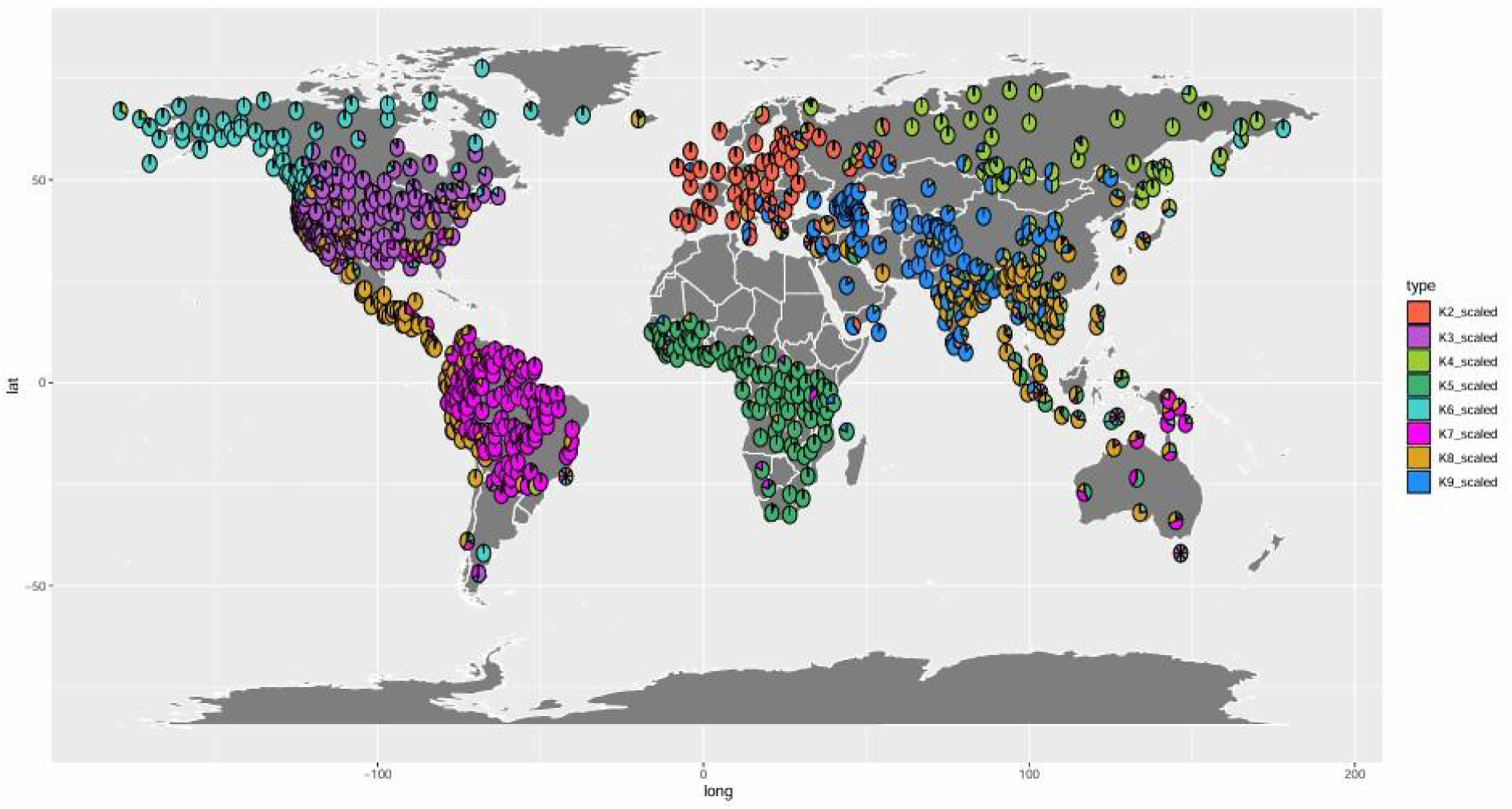
World map displaying ADMIXTURE results for the description of 781 worldwide mythological traditions according to 9 components, with K1 masked.

We further test these two hypotheses by evaluating two models of motif transmission: one of cultural diffusion where the distribution of mythological motifs could be predicted by isolation-by-distance (IBD) alone, and another of demic diffusion according to which mythological motifs would have spread through events of population movement and replacement. Of course, these models only reflect part of the story in and by themselves and represent two extremes of a continuous gradient of mixed mechanisms of transmission which would better fit reality.

### Impact of Ethnolinguistic Barriers

We first use AMOVA^45^ to check for a potential ethnolinguistic structure among the populations of our datasets. For the global dataset (n=73), we obtain ΦST = 0.72 (p-value < 0.001) for genetic distances and ΦST = 0.14 (p-value < 0.001) for mythological ones (Materials and Methods). The first result was much higher than the one obtained by Bortolini et al. (2017)^39^ in their study on folktale diffusion in Eurasia (ΦST = 0.036), yet, it mapped as expected on a PCA space (Fig. S8) of worldwide populations ^46^. We also controlled for the effect of geographical scale on genetic variability by running our analysis on the 25 Eurasian populations we had in common with Bortolini et al. (2017)^39^ and found a ΦST of 0.64 (p-value < 0.001). Their projection in PCA (Fig. S8) also matched that of other studies ^47^ and the Pearson’s product-moment correlation coefficient between their pairwise genetic distances and those obtained by Bortolini and colleagues was of 0.84 (p-value < 0.001). Finally, to test whether our results were due to our choice of ethnolinguistic families, we computed another AMOVA with those used in the paper on folktales, retrieved from Ethnologue (https://www.ethnologue.com), and got ΦST = 0.69 (p-value < 0.001).

Overall, these various tests allow us to be confident in the results of AMOVA.

We did notice a greater variability in the genetic distances obtained from our dataset compared to that of Bortolini and colleagues (Fig. S8), and assumed that the differences in ΦST results could be due to our larger sample size, despite considering a smaller number of genetic markers with respect to their whole genome sequences.

As for ΦST computed with mythological distances, its value was of the same order as that found in previous studies on the spread of cultural elements ^38,39,48^.

### Evaluating Models of Motif Transmission

We explore two models of motif diffusion - one cultural and the other demic - by investigating the relationship between mythological and either geographic or genetic distances. Analyses were also run using distance values corrected for the impact of ethnolinguistic barriers. We then assay the respective fitness of all alternative models by confronting Pearson’s product-moment correlation coefficients, bias-corrected correlations, and partial distance correlations (Materials and methods).

At a worldwide level and using all available motifs, the model associated with the highest correlation values is always that of cultural diffusion hindered by ethnolinguistic barriers (Table 1 “MythemicL∼geographic” and Table 2 “MythemicL∼geographic, geneticL”). As a matter of fact, accounting for the latter improves performance in all cases but that of demic spread rated by partial distance correlations (Fig.2).

**Figure 2.**
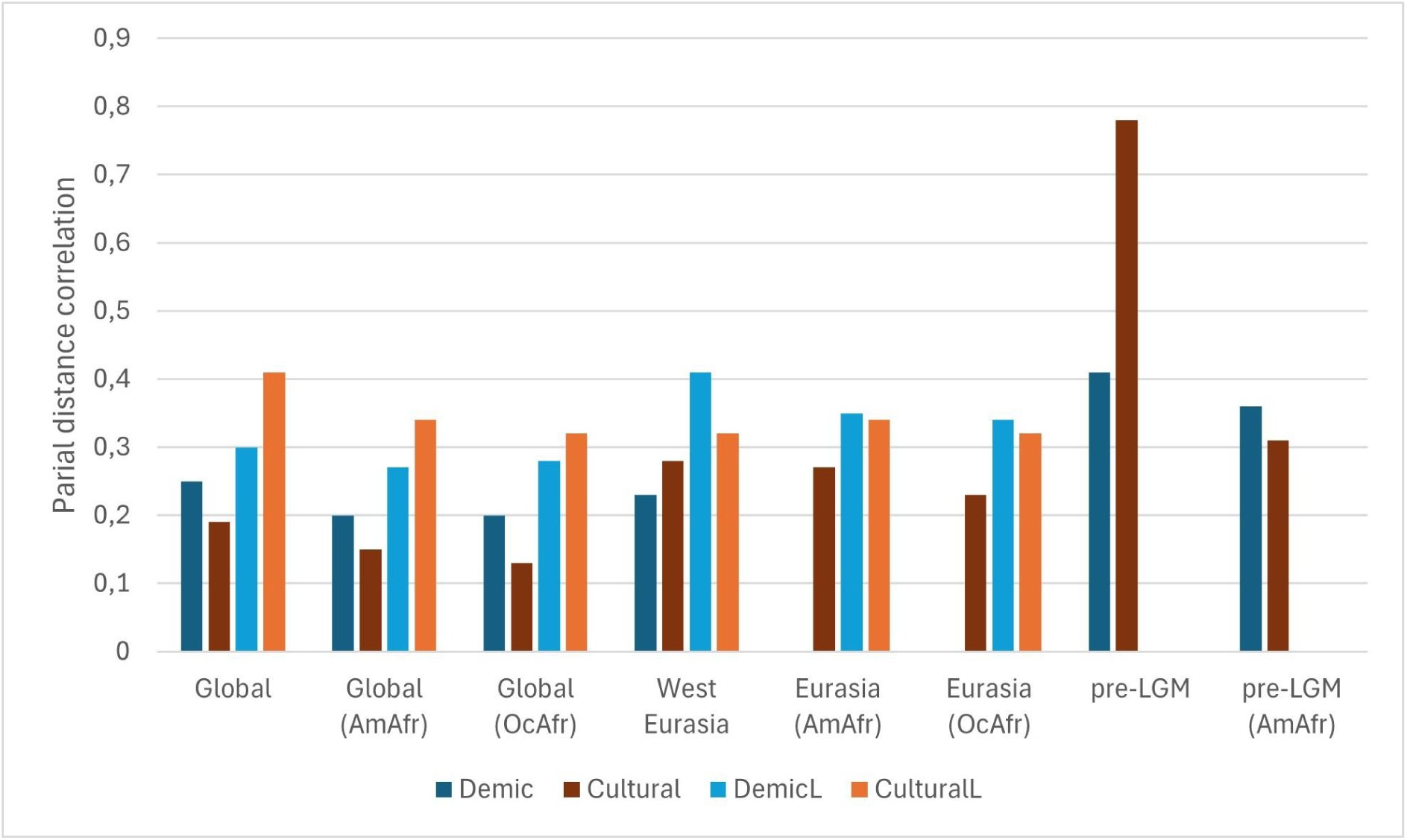
Partial distance correlations associated with the different diffusion models under study (Demic = Mythemic∼genetic, geographic, Cultural = Mythemic∼geographic, genetic, DemicL = MythemicL∼geneticL, geographic, CulturalL = MythemicL∼geographic, geneticL) for the global, West Eurasian, Eurasian, and pre-LGM datasets, either using all motifs available or focusing on AmAfr or OcAfr ones. All p-values are smaller than 0.01.

**Table 1.** Model comparison using the global dataset (n=73) and all available motifs (n=2138) between *demic diffiusion* (mythemic∼genetic), *cultural diffiusion predicted by isolation-by-distance* (mythemic∼geographic), *demic diffiusion biassed by ethnolinguistic barriers* (mythemicL∼geneticL), *cultural diffiusion biassed by ethnolinguistic barriers* (mythemicL∼geographic). The values reported correspond to Pearson’s product-moment correlations (cor) and bias-corrected correlations (b.-c. cor). All p-values are smaller than 0.001.

| Model | cor | b.-c. cor |
| --- | --- | --- |
| Mythemic~genetic | 0.61 | 0.59 |
| Mythemic~geographic | 0.6 | 0.54 |
| Genetic~geographic | 0.77 | 0.73 |
| MythemicL~geneticL | 0.69 | 0.63 |
| MythemicL~geographic | 0.73 | 0.67 |
| GeneticL~geographic | 0.84 | 0.7 |

**Table 2.** Partial distance correlations and their associated p-values for the different models under analysis, using the global dataset (n=73) and all available motifs (n=2138): *demic diffiusion* (mythemic∼genetic, geographic), *cultural diffiusion predicted by isolation-by-distance* (mythemic∼geographic, genetic), *demic diffiusion biassed by ethnolinguistic barriers* (mythemicL∼geneticL, geographic), *cultural diffiusion biassed by ethnolinguistic barriers* (mythemicL∼geographic, geneticL).

| Model | Partial correlation | p-value |
| --- | --- | --- |
| Mythemic~genetic, geographic | 0.35 | < 0.001 |
| Mythemic~geographic, genetic | 0.19 | < 0.001 |
| MythemicL~geneticL, geographic | 0.3 | < 0.001 |
| MythemicL~geographic, geneticL | 0.41 | < 0.001 |

The second best-performing model is globally that of demic diffusion, accounting for ethnolinguistic boundaries (Table 1 “MythemicL∼geneticL”) or not (Table 2 “Mythemic∼genetic, geographic”). However, when we explore model fitness through the computation of Pearson’s product-moment correlation over cumulative geographic distances (Fig. S9), demic processes appear to play a more important role than isolation-by-distance in shaping the mythological landscape below a radius of 5,000km.

Overall, these results suggest that the model explaining most of the current distribution of mythological and folkloric motifs worldwide is that of cultural diffusion impacted by ethnolinguistic barriers. However, partial distance correlations consistently indicate an independent relationship between genetic and mythological distances, which cannot be explained by cultural diffusion alone. The latter even becomes predominant at smaller scales, where genetic and geographic distances correlate less and where their respective effects are thus more easily disentangled (Bortolini et al., 2017).

Focusing on bundles of thematically grouped motifs (Table S8) for which hypothetical routes of demic diffusion were previously proposed in the literature ^11–14,49^, we find consistently lower correlation values between mythological and geographic/genetic distances than we did using the whole dataset. We provide our results and their summarised description in Supplementary materials (Tables S9-S18) and review their implications in the discussion.

Subsequently applying this methodology to West Eurasian populations, for which ancient DNA studies reported substantial genetic re-shaping caused by post-Neolithic demic movements ^50,51^, ethnolinguistic structure appears reduced, though still present (ΦST = 0.37 for genetic distances, ΦST = 0.11 for mythological ones). We thus calculate new parameters for ethnolinguistic barriers and find that Pearson’s product-moment correlation coefficient is maximised when we apply a corrective factor of intensity 0.9 over a range of 1,500 km in the case of genetic distances and over the range of 1,000 km in that of mythological ones.

This time, demic diffusion hindered by ethnolinguistic barriers performs the best out of all tested models (Figure 2), except when evaluated through Pearson’s product-moment correlation, in which case simple demic diffusion performs better (r = 0.69 instead of 0.6) (Tables S19-S20) and does so at any geographic bin (Fig. S10).

We hypothesized that the demic signal detected with both the global and the West Eurasian datasets may refer to prehistoric population movements and we then seeked to reduce confounding effects coming from mostly Eurasian motifs that would have spread through more recent population expansions. For this reason, we built two ensembles. The first comprised motifs shared by at least one African and one South American tradition (AmAfr subset, Table S5), thus theoretically enriching our analysis with a demic signal at least as old as the Last Glacial Maximum, when populations are believed to have crossed the Bering Strait to the Americas (< 23 Kya)^52^. The second one included motifs present in at least one African and one Near Oceanian population (OcAfr subset, Table S6), presumed to have split from other non-African ones at least 30-45 Kya^53,54^.

For both subsets of mythemes, at a worldwide scale, the model performing the best is still that of cultural diffusion limited by ethnolinguistic barriers (Tables S21-24). Interestingly, though, the partial distance correlation between mythological and geographic distances when controlling for the influence of genetic ones is reduced to a greater extent than that between mythological and genetic distances accounting for spatial separation (Figure 2, Tables S22 and S24). The linguistically biased demic model even becomes predominant when we focus exclusively on Eurasian populations (n=56, Materials and Methods), in order to avoid the crossing of large bodies of water from potentially interfering with distances (Tables S25-28). These results suggest that these subsets of mythemes are indeed enriched for information that spread through demic processes, pointing to an ancient (pre-LGM) origin of such a dispersal.

To test this hypothesis, we build genetic distance matrices for 6 time transects (pre-LGM (>20 Kya), 17-9 Kya, 9-6 Kya, 6-4 Kya, 4-2 Kya, present-day) using ancient sequences or present-day ones for regions where, given the broad picture considered here, we could confidently assume population continuity since the Palaeolithic (Materials and Methods, Supplementary Data 2.1-6).

Considering all 2138 motifs, in the case of pre-LGM sequences, we note a statistically significant (p-value < 0.01) relationship between mythological and both genetic and geographic distances with all correlation estimates. (Tables S29 and S30) Interestingly, the demic signal obtained with bias-corrected correlations is stronger than with subsequent time groups until the Bronze Age, after which it reincreases (Fig. 3A, dark blue line). When we account for the effect of geographic distances, on the other hand, the pre-LGM dataset is the only one yielding a statistically significant correlation between mythological and genetic distances (Fig. 3 light blue line, Tables S.32, S34, S36, S38, S40).

**Figure 3.**
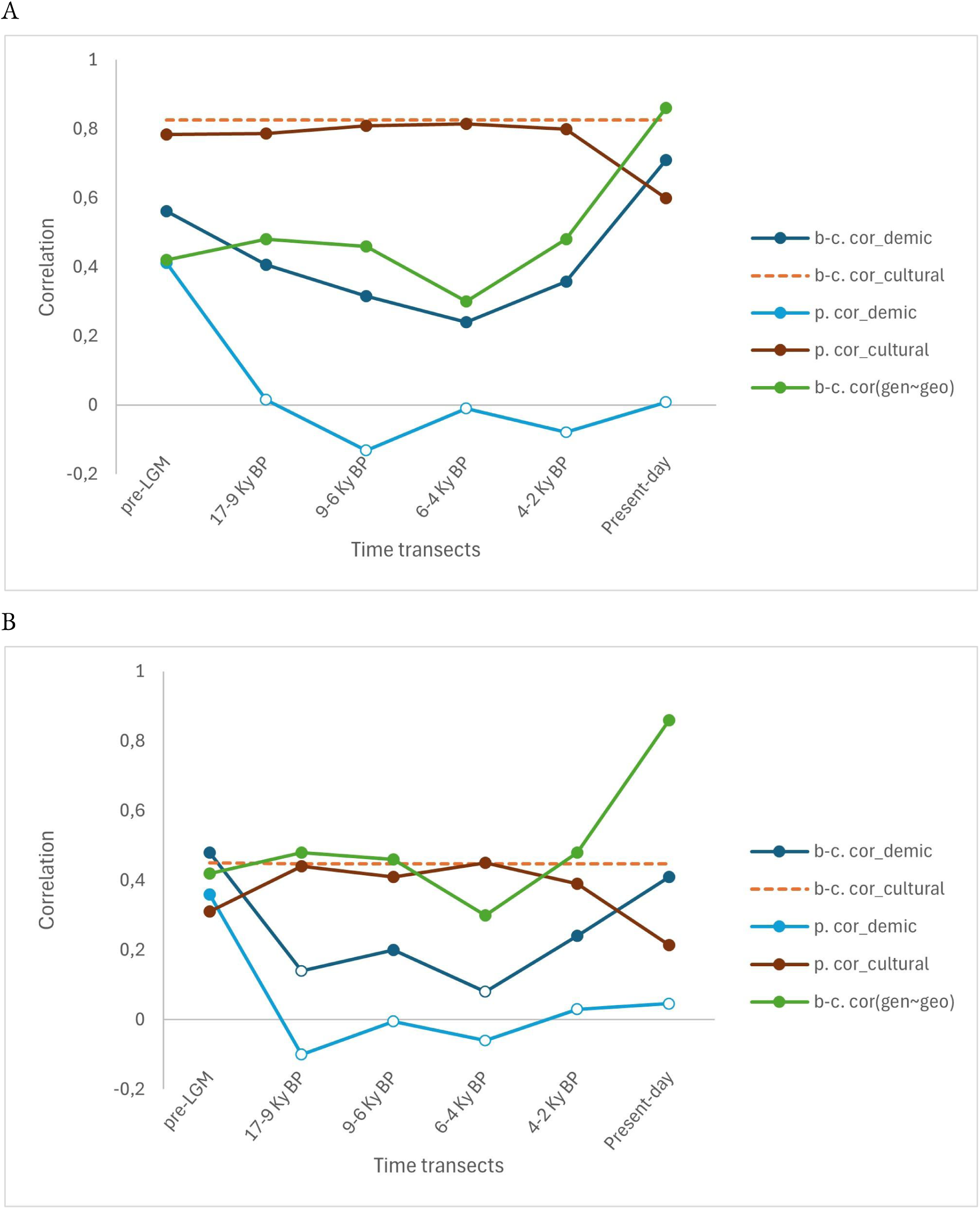
Correlation values over time. We calculated the bias-corrected (b-c. cor) and partial distance correlations (p.cor) between mythological and geographic (cultural model) or genetic (demic model) distances (Materials and Methods) over time for 14 Eurasian populations, either using all motifs present in the dataset (**A**) or using only motifs present in at least one African and one South American populations (Table S5) (**B**). Unfilled dots indicate non-statistically significant relationships (p-value > 0.05).

As with our main dataset, focusing on AmAfr and OcAfr motifs (Table S.5 and S6) in the 6 ensembles increases the relative strength of the demic over the cultural signal (Fig.3B dark blue versus brown lines, Tables S41-S54). After the LGM, we observe a reduction similar to that noted with all motifs and followed by a boost in the Bronze Age. However, this time, we also note a peak during the Neolithic (Fig. 3B) and replicate these general pictures with a smaller dataset including a single European point (Tables S55-S78, Fig. S11). In summary, after accounting for the effect of geography on both mythological and genetic distances, the genetic signal remaining in pre-LGM Eurasia is the only one capable of explaining the present-day distribution of mythological motifs. The relative strength of the demic over the spatial signal also increases when focusing on the AmAfr and OcAfr subsets of motifs (Tables S42 and S44). In other words, of all the genetic signals that cannot be explained by geographic distance alone, the one observed in pre-LGM Eurasia is the only one fitting the observed diffusion of mythemes across the analysed populations. As these results further point toward an enrichment of the AmAfr and OcAfr subsets in motifs older than the LGM, we use a linear regression model (Materials and Methods) to identify those present in both and most strongly associated with a demic signal from Ethiopia, used as a proxy for the exit point Out of Africa. We find 8 motifs corresponding to such criteria and provide their meaning and worldwide distribution maps in Table S.79 and Figure S12.

## Discussion

Before going further into the implications of our main results, it is important to note some limitations of our study. In this paper, we shed light on the time and mechanisms of diffusion of mythological motifs around the world. However, only two models are explored: propagation along a geographical cline and through population movement and replacement. They therefore do not tell the whole story in and by themselves - for example, failing to identify the effects of colonisation and globalisation having taken place over the past 500 years.

Furthermore, many traditions present in Berezkin’s catalogue could not be matched with DNA genotype data and thus had to be excluded from most of our analyses. As for the database itself, biases in its elaboration and discrepancies in the cultural characterisation of traditions of different regions may have reduced the power of our tests (Figures S13 and S14). Its enrichment and standardisation could therefore constitute a significant improvement for prospective studies in comparative mythology.

This notwithstanding, we believe future works on the cultural evolution of our species could benefit from the interdisciplinary approach developed here and inspired by the work of Bortolini et al. (2017)^39^. In particular, the increasing availability of modern and ancient genetic data and advances in computational methods will allow scholars to address ever-more creative and complex questions.

With our work we have compared the worldwide distribution of an extensive collection of mythological motifs with the linguistic and genetic makeup of the human populations that are harbouring them. Our results show that, overall, cultural and demic phenomena concur in explaining the present-day distribution of mythological motifs worldwide. The time depth of these opposite dynamics could be disentangled exploiting available ancient DNA as snapshots of the genetic distribution through space and time.

We first tested subsets of motifs which were hypothesised to have spread via demic diffusion in previous studies^11–14,49^ and, except for the subsets of motifs related to Cosmogony and to a Cosmogonic Dive, dependence between genetic and mythological distances when the effect of geographical ones is taken into account appears almost null and/or statistically insignificant. More in general, the pervasiveness of a mixed signal and the predominance of a spatial cline in some cases should serve as a cautionary tale against unmitigated statements of diffusion through population movement and replacement in comparative mythology studies, as they rather point towards repeated interferences between geography, ethnolingusitic barriers, population movements, and the spread of mythological and folkloric units.

As for our model comparison, on a global scale, motifs appear to have spread predominantly through cultural isolation by distance and partly through demic events. On the other hand, within a spatial range of ca. 5,000 km, our results point to a greater role played by population movements and replacements. Present-day mythological traditions thus appear as the result of various processes of diffusion, in interaction with one another.

The results obtained with motifs present in at least one African and one South American or one Oceanian population, hence presumed to be as old as the settlement of these continents by the first human populations, further suggest distinct histories for different motifs. Notably, in their case, the reduced dependence between mythological and geographic distances at a large scale suggests that we were able to reduce the cultural signal linked to Eurasian interactions having taken place in more recent times. On the other hand, because said dependence wasn’t null, we presume that subsequent transmissions along a geographic cline happened independently in Eurasia and in the Americas over the past 20,000 years and in Eurasia and Oceania since the latter’s settlement.

Finally, within the motifs present in South America, Africa and Oceania, we find that 8 of them present a particularly strong demic signal from Ethiopia, used as a proxy for the exit point Out of Africa (Table S.79). The importance of genetic distance from Ethiopia in explaining their frequency distribution over increasing geographical bins does not mean that they must have exclusively travelled along migration routes, as many kinds of cultural interactions weren’t considered in our analyses. However, we propose that such a signal, combined with the widespread presence of these motifs around the world (Figure S.12) and especially in Africa, South America, and Oceania, supports the hypothesis of their pre-Out-of-Africa origin.

Among them, one refers to the encounter between a woman and a dangerous creature, another to non-human entities superating an obstacle, and a third to food being stolen by women. Then, comes the chase of an animal or object, mistaken for the person or monster of interest, as well as the case in which a small ungulate serves as a trickster. We also find stories about rainbows originating from snakes and water and fish running in tree trunks. Finally, present on all continents, is the tale of the planet Venus being the Moon’s wife.

We then dug further into the time depth of the observed demic signal associated with the spread of mythemes worldwide. On the one hand, our results on West Eurasia and on ancient time groups suggest an important role of Neolithic and Bronze Age population movements in shaping such a spread. On the other hand, simple and partial distance correlations computed at various time transects pointed to the pre-LGM genomic landscape in Eurasia as the only one capable of explaining the demic diffusion of mythemes that presumably originated in Africa and reached Oceania and South America. . Since the analysed pre-LGM landscape originated at least 38 Kya^55^, and in the absence of extensive contacts between Eurasia and Sub-Saharan Africa in palaeolithic times, we conclude that the demic signal we detect could be as old as 60 kya, the time when the expansion Out of Africa of the ancestors of all present-day non-African populations took place. We further provide a shortlist of such myths, which may provide a core set of mythemes that were already known to humankind 60 kya (Table S.79).

Previous studies have found evidence that some stories may have survived into the present day from prehistoric times^30,56^. For example, Nunn and Reid^56^ report that Australian Aboriginal ‘Dreamtime’ myths include specific details about landscapes that vanished with rising sea levels at the end of the LGM over 7,000 years ago, which have since been independently corroborated by physical geographers. Our findings suggest that the earliest surviving myths can be dated back much further into the prehistoric past, constituting the earliest evidence for the existence of storytelling in our species and a unique body of intangible heritage.

## Materials and methods

### Datasets

We retrieved a total of 2138 mythological and folkloric motifs from Yuri Berezkin’s database (Berezkin, 2017), which recorded their presence and absence in 926 traditions in 2018. The latter were also assigned a latitude, a longitude, and up to 3 hierarchical levels of linguistic classification, from broad to narrow scale (for example, Swedish: Indoeuropean - Germanic - Scandinavian). In our analyses, we used Berezkin’s second hierarchical level as a middle ground between linguistic affiliation accuracy and over-specificity, which could have then impacted downstream structure estimations .

To minimise the particularly marked impact of recent contacts with non-African populations attested in North Africa, the Sahel region, and the Western Cape ^57^, we excluded traditions located in Africa and belonging to the Nilo-Saharan and Afrasian linguistic families and kept 88 from the Khoe-San and Niger-Congo ones. They formed the African ensemble used in the rest of our work (Table S1).

Similarly, to reduce the noise deriving from Austronesian interactions ^58^, we elaborated our Near Oceanian ensemble with populations strictly living in Australia and the eastern half of New Guinea, for a total of 17 (Table S2).

We subsequently created five other subsets of populations. The South Asian one (67 populations) covered the region between the Arabian Peninsula and India. The West Eurasian one (73) encompassed Europe and West Asia, while the East Eurasian one (139) did Central and East Asia. Finally, the Americas were divided into a northern (201) and a southern (196) ensemble, with the frontier set in Mexico. Overall, we kept 781 traditions from Berezkin’s original database.

78 of the mythological traditions could be matched with some level of approximation with genetic data retrieved from already published works ^59–81^. In order to prevent confounding signals related to the Out of Africa expansion from interfering with genetic distances (Bortolini et al., 2017)^39^, we kept only the 73 non-African ones in our global dataset and associated each of them with a mythological tradition for which the presence and absence of 2138 motifs were recorded in Berezkin’s database (Supplementary Data 1). We then built a West Eurasian dataset, comprising 21 traditions (Table S3), as well as a Eurasian one including 56 (Table S4).

Besides that, we elaborated 5 ancient datasets. The first one (Supplementary Data 2.1) incorporated populations (n=14) for which we could retrieve sequences antecedent to the Last Glacial Maximum (LGM) or for which we could assume relatively high genetic continuity between present-day and pre-LGM populations as in the cases of Iran, Papuan New Guineans, and Irulas. ^55,82–84^ We chose to include the samples that would best describe the genetic landscape of Palaeolithic Eurasia after the expansion from a population Hub out of Africa at least 38 Kya from which West Eurasians largely descend. ^55^ The other ensembles included the same mythological traditions, this time associated with sequences dated between 17 and 9 Kya (Supplementary Data 2.2), 9 and 6 Kya (Supplementary Data 2.3), 6 and 4 Kya (Supplementary Data 2.4), and 4 and 2 Kya (Supplementary Data 2.5). For these 4 time groups, present-day sequences were used in the cases of Papuan New Guineans, Irulas, and Asiatic Eskimos. All genetic data was retrieved from publicly available sources ^85^ ^86^.

In order to complete this timeline, we built an additional dataset comprising the same mythological traditions, associating them this time with some of the present-day sequences present in our global dataset (Supplementary Data 2.6) Furthermore, we defined two geographic ensembles comprising the 73 non-African populations for which we had genetic data but keeping only mythological motifs found in both Africa and either America (AmAfr subset, Table S5) or Oceania (OcAfr subset, Table S6). To avoid including motifs deriving from later contributions from East Asians to North American groups ^87^, we only used populations from South America for the AmAfr subset (Table S7).

Finally, we performed our analyses on subsets of thematically grouped motifs for which diffusion routes had already been hypothesised in the literature^11–14,49^ (Table S8).

### Admixture between mythological traditions

For the 781 worldwide populations we kept (Materials and Methods), we encoded the presence and absence of mythological motifs as “genetic loci-like” in the form of an ordinary PLINK (.ped) formatted file. We then used the latter to obtain a binary fileset (.bed, .bim., .fam) and run the software ADMIXTURE ^44^. Presence and absence of motifs codified as genetic variants were pruned with a pairwise squared correlation threshold of 0.5 along windows of 50 variants with a shift of 10, and filtered for a minimum frequency of 0.05. For the visualisation of ADMIXTURE results, we settled for 9 components on account of the low cross-validation error associated (Fig. S1).

### Genetic, mythological and geographic distances

Genetic distances between populations included in the global dataset were calculated with Wright’s fixation index (FST) ^35^, computed on PLINK ^88^, using Weir and Cockerham’s method ^89^. On the other hand, because certain populations of the ancient dataset could only be represented by one sequence, genetic distances between them were determined through outgroup-F3 Statistics ^90^, using Mbuti people as an outgroup.

Mythological distance, on the other hand, was obtained with Jaccard’s formula ^91^:

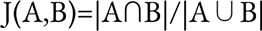

Where A and B are the respective sets of motifs present in two populations, thus allowing us to exclusively infer similarities on the basis of present motifs.

Finally, geographic distance was calculated as pairwise great circle distance, with waypoints in Sinai, the Caucasus, and the Bering Strait to respectively restrict population movements to and from Africa, from South Asia to Eurasia (and vice versa), and into the Americas. The geographic coordinates of each tradition were retrieved from Berezkin’s database.

### AMOVA

We performed AMOVA ^45^ to determine how much genetic and mythological variability between populations was due to their belonging to different ethnolinguistic groups. To do this, we used the ethnolinguistic affiliations listed in Berezkin’s database ^5^ and the function amova available in the package pegas ^92^ in R. P-values, assessing the significance of results, were estimated on the basis of 1,000 iterations.

### Accounting for Ethnolinguistic Barriers

In order to account for ethnolinguistic barriers, which can constrain the diffusion of cultural traits and genes ^38,39^, we applied the methodology introduced by Bortolini et al. (2017)^39^. An intensity factor f and a geographical radius of impact r were thus independently computed for genetic and mythological distances. The correlation between the latter and geographic distance was estimated over increasing spatial bins (r), with and without the correction calculated as:

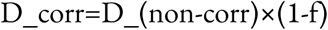

Where f could take values from 0 to 1, with increments of 0.1.

In the end, the chosen pair of r and f was that which maximised said correlation and linguistically corrected distances were obtained by applying the above equation to all pairs of populations falling within a distance < r from one another. We found that the correlation between geographic and genetic/mythological distances is maximised when we respectively correct the latter with a factor of intensity of 0.3 and 0.2 under a geographic range of 10,000 and 5,000 km.

### Model comparison

The fitness of each model (cultural or demic diffusion, with or without ethnolinguistic barriers) was determined by Pearson’s product-moment correlation coefficient with the function cor.test in R, a bias-corrected adaptation of distance correlation based on a nonparametric t-test of multivariate independence in high dimension, using the function dcorT.test from the package energy in R ^93^, and partial distance correlations, computed with the function pdcor.test in the package energy in R.

In order to implement the latter two, distances had to be Euclidean.^93,94^ Mythological and genetic ones were thus transformed into their exact Euclidean version through the classical multidimensional scaling of their matrices, done with the function cmdscale in R, on which was then used the function dist in R.^95^ As for geographic distances, we used the function dist.mat of the package REAT in R, which computes Euclidean distances between sets of spatial coordinates.

Pearson’s product-moment correlation coefficients were also computed over cumulative geographical distance bins to allow for different outcomes depending on the scale of analysis.

### Genetic Effect Sizes of Mythological Motifs

We calculated the geographic distance between all 781 populations with a mythological tradition present in Berezkin’s database and Ethiopia, used as a proxy for the exit point Out of Africa, in the same way as described above. We then min-max normalised them and classified them into distance bins of 0.05 from 0 to 0.5 and of 0.1 from there to 1 to account for the reduced number of included populations at a larger scale. To investigate the strength of the demic signal associated with each motif, we then described their frequency in each bin as the linear combination of the mean min-max normalised genetic and geographic distances from Ethiopia in said bin, using data from the 78 populations of our global dataset. The model was computed with the function lm in R and the effect size of each motif’s genetic coefficient with the function omega_squared from the package effectsize in R. We gauged the value of the latter according to Cohen’s criterion. ^96^

## Data Availability Statement

Mythological data used for this work was made available by Yuri Berezkin and is available as Supplementary Data 1. Genetic data was obtained from publicly available sources at the Institute of Genomics of the University of Tartu https://ebc.ee/free_data/ and at the Allen Ancient DNA Resource https://dataverse.harvard.edu/dataset.xhtml?persistentId=doi:10.7910/DVN/FFIDCW

## Acknowledgements

The authors have no conflict of interest to report. Luca Pagani would like to thank Prof. Richard Villems for introducing him to the work of Michael Witzel. LP was funded by the Italian Ministry of Research PRIN N. P2022MZCNX.

## References

1. Berezkin, Y. E. Cosmic hunt: Variants of Siberian-north American myth. Folk. Electron. J. Folk. 31, 79–100 (2005).

2. Berezkin, Y. E. Earth-diver emergence from under earth: Cosmogonic tales evidence favor heterogenic origins American Indians. Ethnology Anthropology Eurasia 32, 110–123 (2007).

3. Berezkin, Y. E. Dwarfs Cranes: Baltic Finnish Mythologies Eurasian American Perspective (70 Years after Yriö Toivonen). Folklore: Electronic Journal Folklore 36, 75–96 (2007).

4. Berezkin, Y. E. Spread of folklore motifs as a proxy for information exchange: contact zones and borderlines in Eurasia. Trames J. Humanit. Soc. Sci. 19, 3 (2015).

5. Berezkin, Y. E. Peopling of the new world from data on distributions of folklore motifs. in Maths Meets Myths: Quantitative Approaches to Ancient Narratives 71–89 (Springer International Publishing, Cham, 2017).

6. Berezkin, Y. E. Stratigraphy of cultural interaction in Eurasia based on computing of folklore motifs. Trames J. Humanit. Soc. Sci. 20, 217 (2016).

7. d’Huy, J. Cosmic Hunt Berber Sky: Phylogenetic Reconstruction Palaeolithic Mythology. Les Cahiers de 93–106 (2013).

8. d’Huy, J. Phylogenetic Reconstruction Prehistoric Tale. Nouvelle Mythologie Comparée/New Comparative Mythology. 3–18 (2013).

9. d’Huy, J. Un récit de plongeon cosmogonique au Paléolithique supérieur ? Préhistoire du Sud-Ouest 25, 109–117 (2017).

10. d’Huy, J. Lascaux, les Pléiades et la Voie lactée: à propos d’une hypothèse en archéoastronomie. Mythologie française 19–22 (2017).

11. d’Huy, J. Matriarchy prehistory: statistical method testing old theory. Les Cahiers de l’AARS 19, 159–170 (2017).

12. d’Huy, J. Coda : La symphonie du premier plongeon. Préhistoire du Sud-Ouest 26, 179–190 (2018).

13. d’Huy, J. « Mort, où est ta victoire» Reconstruction statistique des premières croyances de l’humanité sur la mort. Paléo 182–195 (2020).

14. d’Huy, J. & Berezkin, Y. E. How Did First Humans Perceive Starry Night? On the Pleiades. Retrospective Methods Network Newsletter 100–122 (2017).

15. d’Huy, J., et al. Little Statisticians Forest Tales: Towards New Comparative Mythology. Fabula 44–63 (2023).

16. Hafstein, V. Biological metaphors in folklore theory: an essay in the history of ideas. Arv 57, 7–32 (2001).

17. Jacobs, J. European Folk and Fairy Tales. (General Books, 2010).

18. Jung, C. Archetypes Collective Unconscious, Collected Works. vol. 9 (Princeton University Press, 1959).

19. Konstan, D. Comparative Methods in Mythology. Arethusa 9, (1986).

20. Korotayev, A. & Khaltourina, D. Glubokaja istoričeskaja rekonstrukcija. Glubokaja istoričeskaja rekonstrukcija (2011).

21. Levi-Strauss, C. The structural study of myth. J. Am. Folk. 68, 428 (1955).

22. Lévi-Strauss, C. L’homme nu. (Librairie Plon, 1971).

23. Lévi-Strauss, C. De Grées ou de force ? Homme 7–18 (2002).

24. Thuillard, M., Le Quellec, J.-L., d’Huy, J. & Berezkin, Y. E. A large-scale study of world myths. Trames J. Humanit. Soc. Sci. 22, 407 (2018).

25. Uther, H.-J. Types International Folktales: Classification Bibliography. Based System Antti Aarne Stith Thompson. (Helsinki: Suomalainen Tiedeakatemia). (2004).

26. Michael Witzel, E. J. The Origins of the World’s Mythologies. (Oxford University Press, New York, NY, 2012).

27. Bhattacharya, D. Folktales Folder Human Mind: Analytical Overview. Folktales Folder Human Mind: Analytical Overview. JHSSS 3,.

28. Lévi-Strauss, C. La potière jalouse. (1985).

29. Thompson, S. Types Folktale: Classification Bibliography. Antti Aarne’s Verzeichnis Der Märchentypen. (Suomalainen Tiedeakatemia, Helsinki, 1928).

30. Da Silva, S. G. & Tehrani, J. J. Comparative phylogenetic analyses uncover ancient roots Indo-European folktales. R. Soc. Open Sci 3, (2016).

31. d’Huy, J. Un ours dans les étoiles, recherche phylogénétique sur un mythe préhistorique. Préhistoire du Sud-Ouest 20, 91–106 (2012).

32. Thuillard, M. & Le Quellec, J. L. A phylogenetic interpretation canonical formula myths by Levi-Strauss. CAES 2, (2017).

33. Tehrani, J. J. The phylogeny of Little Red Riding Hood. PLoS One 8, e78871 (2013).

34. Sakamoto Martini, G., Kendal, J. & Tehrani, J. J. Cinderella’s Family Tree. Phylomemetic Case Study ATU 510/511. Fabula 64, 7–30 (2023).

35. Wright, S. Genetical structure of populations. Nature 166, 247–249 (1950).

36. Rzeszutek, T., Savage, P. E. & Brown, S. The structure of cross-cultural musical diversity. Proc. Biol. Sci. 279, 1606–1612 (2012).

37. Bell, A. V., Richerson, P. J. & McElreath, R. Culture rather than genes provides greater scope for the evolution of large-scale human prosociality. Proc. Natl. Acad. Sci. U. S. A. 106, 17671–17674 (2009).

38. Ross, R. M., Greenhill, S. J. & Atkinson, Q. D. Population structure and cultural geography of a folktale in Europe. Proc. Biol. Sci. 280, 20123065 (2013).

39. Bortolini, E. et al. Inferring patterns of folktale diffusion using genomic data. Proc. Natl. Acad. Sci. U. S. A. 114, 9140–9145 (2017).

40. Lycett, S. J. Cultural evolutionary approaches to artifact variation over time and space: basis, progress, and prospects. J. Archaeol. Sci. 56, 21–31 (2015).

41. Fort, J. Synthesis between demic and cultural diffusion in the Neolithic transition in Europe. Proc. Natl. Acad. Sci. U. S. A. 109, 18669–18673 (2012).

42. Barbieri, C. et al. A global analysis of matches and mismatches between human genetic and linguistic histories. Proc. Natl. Acad. Sci. U. S. A. 119, e2122084119 (2022).

43. Harris, E. E. Demic and cultural diffusion in prehistoric Europe in the age of ancient genomes. Evol. Anthropol. 26, 228–241 (2017).

44. Alexander, D. H., Novembre, J. & Lange, K. Fast model-based estimation of ancestry in unrelated individuals. Genome Res. 19, 1655–1664 (2009).

45. Excoffier, L., Smouse, P. E. & Quattro, J. M. Analysis of molecular variance inferred from metric distances among DNA haplotypes: application to human mitochondrial DNA restriction data. Genetics 131, 479–491 (1992).

46. Almarri, M. et al. Xue, Y. Population Structure, Stratification, Introgression Human Structural Variation. Cell 182, 189–199 (2020).

47. Allentoft, M. E. et al. Population genomics of post-glacial western Eurasia. Nature 625, 301–311 (2024).

48. Shennan, S. J., Crema, E. R. & Kerig, T. Isolation-by-distance, homophily, core vs. package cultural evolution models Neolithic Europe. Evolution Human Behavior 103–109 (2015).

49. d’Huy, J. Cosmogonies. (La Découverte, 2020).

50. Haak, W. et al. Massive migration from the steppe was a source for Indo-European languages in Europe. Nature 522, 207–211 (2015).

51. Lazaridis, I. et al. Genomic insights into the origin of farming in the ancient Near East. Nature 536, 419–424 (2016).

52. Raghavan, M. et al. Genomic evidence for the Pleistocene and recent population history of Native Americans. Science 349, aab3884–aab3884 (2015).

53. Choin, J. et al. Genomic insights into population history and biological adaptation in Oceania. Nature 592, 583–589 (2021).

54. O’Connell, J. F., et al. When did Homo sapiens first reach Southeast Asia and Sahul? Proc. Natl. Acad. Sci. U. S. A. 115, 8482–8490 (2018).

55. Vallini, L. et al. Genetics and material culture support repeated expansions into paleolithic Eurasia from a population Hub out of Africa. Genome Biol. Evol. 14, (2022).

56. Nunn, P. D. & Reid, N. J. Aboriginal Memories Inundation Australian Coast Dating from More than 7,000 Years Ago. Australian Geographer (2015).

57. Pfennig, A., Petersen, L. N., Kachambwa, P. & Lachance, J. Evolutionary genetics and admixture in African populations. Genome Biol. Evol. 15, (2023).

58. Oliveira, S. et al. Ancient genomes from the last three millennia support multiple human dispersals into Wallacea. Nat. Ecol. Evol. 6, 1024–1034 (2022).

59. Tätte, K. et al. The genetic legacy of continental scale admixture in Indian Austroasiatic speakers. Sci. Rep. 9, 3818 (2019).

60. Ongaro, L. et al. The genomic impact of European colonization of the Americas. Curr. Biol. 29, 3974–3986.e4 (2019).

61. Grugni, V. et al. Analysis human Y-chromosome: haplogroup Q characterizes ancient population movements Eurasia Americas. BMC Biol 17, (2019).

62. Bryc, K. et al. Genome-wide patterns of population structure and admixture in West Africans and African Americans. Proc. Natl. Acad. Sci. U. S. A. 107, 786–791 (2010).

63. Bryc, K., Durand, E. Y., Macpherson, J. M., Reich, D. & Mountain, J. L. The genetic ancestry of African Americans, Latinos, and European Americans across the United States. Am. J. Hum. Genet. 96, 37–53 (2015).

64. Cardona, A. et al. Genome-wide analysis of cold adaptation in indigenous Siberian populations. PLoS One 9, e98076 (2014).

65. Hoffecker, J. F., Elias, S. A., O’Rourke, D. H., Scott, G. R. & Bigelow, N. H. Beringia and the global dispersal of modern humans. Evol. Anthropol. 25, 64–78 (2016).

66. Meade, T. History Modern Latin America: 1800 Present. (John Wiley & Sons, 2011).

67. Migliano, A. B. et al. Evolution of the pygmy phenotype: Evidence of positive selection from genome-wide scans in African, Asian, and Melanesian pygmies. Hum. Biol. 85, 251 (2013).

68. Mörseburg & Pagani, L. R. Multi-layered population structe Island Southeast Asians. Eur J Hum Genet 24, 1605–1611 (2016).

69. Pagani, L. et al. Genomic analyses inform on migration events during the peopling of Eurasia. Nature 538, 238–242 (2016).

70. Moreno-Estrada, A. et al. Human genetics. The genetics of Mexico recapitulates Native American substructure and affects biomedical traits. Science 344, 1280–1285 (2014).

71. Li, J. Z. et al. Worldwide human relationships inferred from genome-wide patterns of variation. Science 319, 1100–1104 (2008).

72. Kehdy, F. S. G. et al. Origin and dynamics of admixture in Brazilians and its effect on the pattern of deleterious mutations. Proc. Natl. Acad. Sci. U. S. A. 112, 8696–8701 (2015).

73. Clemente, F. J. et al. A selective sweep on a deleterious mutation in CPT1A in arctic populations. Am. J. Hum. Genet. 95, 584–589 (2014).

74. Busby, G. B. J. et al. The role of recent admixture in forming the contemporary west Eurasian genomic landscape. Curr. Biol. 25, 2878 (2015).

75. Goldberg, A. & Rosenberg, N. A. Beyond 2/3 and 1/3: The complex signatures of sex-biased admixture on the X chromosome. Genetics 201, 263–279 (2015).

76. Tamm, E. et al. Beringian standstill and spread of Native American founders. PLoS One 2, e829 (2007).

77. Chacón-Duque, J.-C. et al. Latin Americans show wide-spread Converso ancestry and imprint of local Native ancestry on physical appearance. Nat. Commun. 9, 5388 (2018).

78. Moreno-Mayar, J. V. et al. Terminal Pleistocene Alaskan genome reveals first founding population of Native Americans. Nature 553, 203–207 (2018).

79. Moreno-Mayar, J. V. et al. Early human dispersals within the Americas. Science 362, eaav2621 (2018).

80. Henn, B. M. et al. Hunter-gatherer genomic diversity suggests a southern African origin for modern humans. Proc. Natl. Acad. Sci. U. S. A. 108, 5154–5162 (2011).

81. The International HapMap 3 Consortium. Integrating common and rare genetic variation in diverse human populations. Nature 467, 52–58 (2010).

82. Yelmen, B. et al. Ancestry-specific analyses reveal differential demographic histories and opposite selective pressures in modern South Asian populations. Mol. Biol. Evol. 36, 1628–1642 (2019).

83. Vallini, L. et al. The Persian plateau served as hub for Homo sapiens after the main out of Africa dispersal. Nat. Commun. 15, 1882 (2024).

84. Guarino-Vignon, P., Marchi, N., Bendezu-Sarmiento, J., Heyer, E. & Bon, C. Genetic continuity of Indo-Iranian speakers since the Iron Age in southern Central Asia. Sci. Rep. 12, 733 (2022).

85. Mallick, S. & Reich, D. The Allen Ancient DNA Resource (AADR): A curated compendium of ancient human genomes. Harvard Dataverse 10.7910/DVN/FFIDCW (2023).

86. Mallick, S. et al. The Allen Ancient DNA Resource (AADR) a curated compendium of ancient human genomes. Sci. Data 11, 182 (2024).

87. Willerslev, E. & Meltzer, D. J. Peopling of the Americas as inferred from ancient genomics. Nature 594, 356–364 (2021).

88. Purcell, S. et al. PLINK: toolset whole-genome association population-based linkage analysis. American Journal Human Genetics 559–575 (2007).

89. Weir, B. S. & Cockerham, C. C. Estimating F-statistics for the analysis of population structure. Evolution 38, 1358 (1984).

90. Patterson, N. et al. Ancient admixture in human history. Genetics 192, 1065–1093 (2012).

91. Jaccard, P. Étude comparative de la distribution florale dans une portion des Alpes et des Jura. Bulletin de la Société vaudoise des sciences naturelles 37, 547–579 (1901).

92. Paradis, E. pegas: an R package for population genetics with an integrated-modular approach. Bioinformatics 26, 419–420 (2010).

93. Szekely, G. & Rizzo, M. distance correlation t-test independence high dimension. Journal Multivariate Analysis 117, 193–213 (2013).

94. Székely, G. J., Rizzo, M. L. & Bakirov, N. K. Measuring and testing dependence by correlation of distances. Ann. Stat. 35, 2769–2794 (2007).

95. Székely, G. J. & Rizzo, M. L. Partial distance correlation with methods for dissimilarities. Ann. Stat. 42, 2382–2412 (2014).

96. Cohen, J. Statistical Power Analysis for the Behavioral Sciences. (Elsevier Science, 2013).

