## Supplementary Information for "Worldwide patterns in mythology echo the human expansion out of Africa"

**Table S1.** African populations present in Berezkin's database and used to form our African ensemble.

| |
|---|
| Bushmen; Baule, Nzema, Agni; Vai, Kone; Zande,Nzakara; Khoikhoi; Akan,Ashanti,Akwapim,Tvi; Kpelle, Kono; Mbum,Mundang; Xhosa; Fon; Mende,Bandi,Loma; Hadza; Zulu,Swasi; Ewe; Manden,Bamana,Malinke,Diula; Sandawe; Sotho, Tswana; Akposo,Akebu,Santrokofi; Susu; Makaa, Kaka; Tonga, Ndebele; Northern Gur (Oti-Volta); Soninke; Fang,Bube,Bulu,Tanga; Shona; Southern Gur; Bozo,Sorko; Mpongwe,Nkomi,Mindumu; Bemba,Kaonde,Lamba; Lobi; Bobo; Aka,Baka a.o.Western Pygmies; Tsonga,Soli,Sala,Lenje; Senufo; Dan,Guro,Mano,Sapa,Ngere; Yambasa, Banen;; Ila; Dogon; Kalenjin,Nande,Arusha; Tiv,Jukun,Bete,Wute; Malawi; Fulbe,Wolof,Serer; Mangbetu,Lugbara,Madi; Efik, Ibibio, Ikom; Nyungwe; Manjak, Balant, Papel,Felupe; Ngonde,Safwa.Mkulwe,Kinga; Yoruba,Nupe,Bini; Yao, Macua; Tenda, Biafada, Nalu; Konde,Matumbi,Pangwe; Ijaw; Herrero; Temne; Swahili; Duala,Basa,Kwiri; Lozi,Rotse,Lui,Subia; Limba; Comoros Islands; Rwanda,Shi,Rundi; Mbundu, Owambo; Gola; Banda,Gbaya,Ngbandi,Manja; Ganda,Nyoro; Chokwe,Luchazi; Kissi; Kikuyu; Niamwesi,Sumbwa; Lunda; Kru; Chagga, Digo; Nyaturu,Isanzu,Nilamba; Kongo; Luba,Bena,Tabwa; Gogo,Zaramo,Kaguru; Sukuma,Gusii,Kumbi; Sakata; Lega, Bangubangu; Yaka; Fipa,Bende,Iramba; Kuba,Dengese; Songe; Boa,Komo,Nyanga; Kamba; Lingala; Soko; Mongo,Tetela,Kuba,Nkundu; Kete,Luba-Kasai,Kaniok |

**Table S2.** Australian and eastern New Guinean populations used to constitute the "Oceanian ensemble".

| |
|---|
| Bunak; Fataluku; Halmahera; Tasmania; SE Australia; Arnham Land; Kimberley Plateau; Trans New Guinea East; Torricelli Papuans; Sepik-Ramu Papuans; Torres Strait Islands; North New Guinea Melanesians; SE New Guinea Melanesians; Queensland; Central Australia; Southern Australia; Western Australia |

**Table S3.** Populations included in the West Eurasian dataset, referred to with the names given in the "Genetic population" column of Table 3.

| |
|---|
| Abkhasians; French; Poles; Adygei; Germans; Portuguese; Balkars; Ingrians; Russians; Belarusians; Italians; Saami_merged; Croats; Karelian; Swedes; Estonians; Latvians; Ukranians; Finns; Lithuanians; Vepsas |

**Table S4.** Populations included in the Eurasian dataset, referred to with the names given in the "Genetic population" column of Table 3.

| |
|---|
| Altaians; Balkars; Bashkirs; Batak; Belarusians; Borneo_merged; Burmese; Buryats; Chukchis; Chuvash; Croats; Eskimo; Estonians; Evenkis_merged; Evens; Finns; French; Germans; Han; Hantis; Igorot; Ingrians; Italians; Japanese; Karelian; Kazakhs; Komis; Koryaks_merged; Kyrgyzians; Latvians; Lezgins; Lithuanians; Malay; Mansis; Maris; Mongolians_merged; Mordovians; Nenets_merged; Nganasans; North_Ossetians; Pamiris; Poles; Portugese; Russians; Saami_merged; Shors; Swedes; Tajiks; Turkmens; Tuvinians; Udmurts; Ukranians; Vepsas; Vietnamese; Yakuts |

**Table S5.** AmAfr motifs (n=346)

| | | | | | | | | | | | |
|---|---|---|---|---|---|---|---|---|---|---|---|
| a1_1 | a2a_1 | a2d_1 | a3_1 | a4_1 | a5_1 | a6_1 | a11a_1 | a12_1 | a12c_1 | a12d_1 | a14_1 |
| a15_1 | a16_1 | a17_1 | a21_1 | a24_1 | a32_2 | a32a_2 | a32e_2 | a35_2 | a36_4 | a37_1 | a38_1 |
| b1_3 | b2a_3 | b2d_3 | b2e_3 | b3a_3 | b3b_3 | b3e_3 | b7b_3 | b9_3 | b13a_10 | b14_3 | b15_3 |
| b17_3 | b18_3 | b30b_7 | b37_7 | b38_7 | b40_7 | b40a_7 | b41_7 | b42_2 | b42k_2 | b42n_2 | b77_3 |
| b86_3 | b98_7 | b102_3 | c1_3 | c5a_3 | c5b_3 | c6f_10 | c10_7 | c18_3 | c19_3 | c23_3 | d1b_3 |

| | | | | | | | | | | | |
|---|---|---|---|---|---|---|---|---|---|---|---|
| d4a_3 | d4j_7 | d4l_3 | d4o_11 | d5_3 | d8_7 | d9_7 | d11_6 | d12_6 | d13hh_8 | e1b_5 | e5a_5 |
| e5b_5 | e5c_5 | e8_5 | e9_10 | e9c_5 | e10_5 | e11_10 | e14_6 | e24_5 | e32_5 | e35_5 | f1_5 |
| f2_5 | f4_5 | f7_5 | f8_5 | f9_5 | f9a_5 | f17_5 | f18b_5 | f22_5 | f27_5 | f30_5 | f35_10 |
| f38_5 | f39_5 | f40a_5 | f40b_5 | f40c_10 | f42_5 | f43a_5 | f44_5 | f45_5 | f50_5 | f55_5 | f58_11 |
| f61_11 | f64_11 | f65_11 | f65a_10 | f73_5 | f76_5 | f80_5 | f80a_5 | f86_10 | f99_5 | g6_6 | g8b_10 |
| g9_10 | g13a_6 | g13b_6 | g23_10 | g28_6 | h1a_4 | h1c_4 | h2_4 | h3_4 | h4_4 | h4a_4 | h5_4 |
| h6a_4 | h6bb_4 | h7_8 | h8_4 | h9_4 | h10_4 | h11_8 | h12_8 | h18_7 | h20a_7 | h21_10 | h24_10 |
| h24a_2 | h24c_4 | h27_8 | h28_8 | h34a_4 | h34g_4 | h36_11 | h36a_4 | h36c_4 | h36ff_4 | h36hh_4 | |
| h39_7 | h41_4 | h47_3 | i1_3 | i2_3 | i3_3 | i4a_10 | i5_3 | i5a_3 | i7_3 | i8f_3 | i8g_3 |
| i9_3 | i10a_3 | i11_3 | i13a_8 | i14_8 | i14a_8 | i16_5 | i20_8 | i26_8 | i27_8 | i28_3 | i32_5 |
| i33_8 | i35_3 | i39_3 | i40_3 | i41_3 | i41a_3 | i42_3 | i43a_3 | i43b_3 | i44_3 | i45a_8 | i45b_8 |
| i45c_8 | i46_3 | i47_3 | i55_2 | i62_2 | i65_2 | i66_2 | i69_2 | i72_2 | i73_2 | i74_2 | i74a_2 |
| i78_3 | i82a_2 | i82b_2 | i82c_2 | i100_2 | i100a_2 | i100b_2 | i103_2 | i104_2 | i108_2 | i115a_2 | i117_7 |
| j4_10 | j7_10 | j12d_10 | j15_10 | j25_10 | j26_10 | j46_10 | j47_10 | j56_10 | j56a_10 | j58_10 | k1f_10 |
| k2_10 | k4_10 | k8a_10 | k8c_10 | k8d_10 | k8e_11 | k11a_7 | k12_10 | k15a_10 | k18_10 | k22_3 | k23_10 |
| k24_10 | k24a_10 | k25_10 | k25a1_10 | k27e_10 | k27i_10 | k27n_10 | k27n1_10 | k27n3a_10 | k27n3b_10 | k27p_10 | |
| k27s_10 | k28_10 | k29a_10 | k32_10 | k33_10 | k35_10 | k44_10 | k44a_10 | k55_10 | k56b_10 | k60b_10 | k61b_10 |
| k75_10 | k77a_10 | k79_10 | k86_10 | L4_10 | L5c_8 | L6_8 | L7_10 | L9a_8 | L13_10 | L14_8 | L15a_8 |
| L17a_8 | L17b_8 | L18_8 | L19b_8 | L21_10 | L27_10 | L32_8 | L33_8 | L34_10 | L40_10 | L41_10 | L41a_10 |
| L42_10 | L42b_10 | L45_11 | L52_10 | L53_10 | L55_10 | L56_10 | L61_10 | L63_10 | L64_8 | L65_10 | L68_10 |
| L70_11 | L72_10 | L73_10 | L85_8 | L92_10 | L103_10 | L110_10 | m2_10 | m3_10 | m5_11 | m8_10 | m8a_10 |
| m11c_10 | m12_10 | m21_10 | m23_11 | m25_11 | m28_11 | m29b_9 | m29g_9 | m29i_9 | m29k_9 | m29o_9 | m29u_9 |
| m29v_9 | m30_11 | m33_11 | m38_11 | m40_11 | m42_11 | m44b_11 | m47_10 | m50_11 | m51_11 | m57a_8 | m57b_8 |
| m60b_11 | m62a_11 | m63_10 | m72_10 | m81_10 | m87_11 | m89_11 | m102_11 | m115_11 | m183_11 | m185_11 | |
| m185a_11 | | | | | | | | | | | |

**Table S6.** OcAfr motifs (n=225)

| | | | | | | | | | | | |
|---|---|---|---|---|---|---|---|---|---|---|---|
| a3_1 | a4_1 | a5_1 | a11a_1 | a12_1 | a12c_1 | a14_1 | a17_1 | a21_1 | a24_1 | a32_2 | a32e_2 |
| a35_2 | a36_4 | a38_1 | a41_1 | b2a_3 | b2d_3 | b2e_3 | b3_3 | b3a_3 | b3b_3 | b3e_3 | b7b_3 |
| b9_3 | b13a_10 | b14_3 | b30b_7 | b37_7 | b38_7 | b41_7 | b42_2 | b42n_2 | b77_3 | c5a_3 | c6f_10 |
| d4a_3 | d4e1_7 | d4l_3 | d5_3 | d9_7 | d11_6 | d12_6 | d13hh_8 | e5a_5 | e5b_5 | e5c_5 | e8_5 |
| e9_10 | e10_5 | e11_10 | e32_5 | e35_5 | f1_5 | f2_5 | f4_5 | f7_5 | f8_5 | f9_5 | f9a_5 |
| f18b_5 | f27_5 | f30_5 | f35_10 | f38_5 | f39_5 | f40a_5 | f40b_5 | f40c_10 | f42_5 | f43a_5 | f44_5 |
| f45_5 | f55_5 | f58_11 | f64_11 | f65_11 | f80_5 | f80a_5 | f86_10 | f97_5 | f99_5 | g6_6 | g8_10 |
| g8b_10 | g13a_6 | g23_10 | g28_6 | h1c_4 | h4_4 | h4a_4 | h5_4 | h7_8 | h9_4 | h10_4 | h12_8 |
| h12b_8 | h18_7 | h20a_7 | h24_10 | h27_8 | h34a_4 | h36ff_4 | h36hh_4 | h41_4 | i1_3 | i3_3 | i7_3 |
| i8f_3 | i8g_3 | i13a_8 | i14_8 | i14a_8 | i16_5 | i20_8 | i27_8 | i28_3 | i32_5 | i39_3 | i41_3 |
| i42_3 | i43a_3 | i43b_3 | i44_3 | i45b_8 | i47_3 | i62_2 | i65_2 | i66_2 | i72_2 | i74_2 | i82a_2 |
| i82b_2 | i82c_2 | i100_2 | i100b_2 | i108_2 | i115a_2 | i116_2 | i119_8 | j4_10 | j15_10 | j23_10 | j25_10 |
| j26_10 | j46_10 | j47_10 | j58_10 | k1f_10 | k2_10 | k4_10 | k8a_10 | k8c_10 | k11a_7 | k12_10 | k15a_10 |
| k24_10 | k25_10 | k25a1_10 | k27hh_10 | k27n1_10 | k27n3b_10 | k28_10 | k29a_10 | k32_10 | k33_10 | k33b_10 | |
| k37_10 | k49_10 | k56b_10 | k74_10 | k75_10 | k76_10 | k77a_10 | k83_10 | k86_10 | L4_10 | L5c_8 | L6_8 |
| L7_10 | L15a_8 | L15d_8 | L17a_8 | L19b_8 | L21_10 | L40_10 | L41_10 | L42_10 | L50_10 | L52_10 | L53_10 |
| L55_10 | L61_10 | L64_8 | L65_10 | L70_11 | L81_10 | L85_8 | L85a_10 | L106_10 | L108_10 | L110_10 | |
| m3_10 | m3a_11 | m8_10 | m8a_10 | m12_10 | m23_11 | m29k_9 | m29o_9 | m29u_9 | m29v_9 | m33_11 | m42_11 |
| m44b_11 | m50_11 | m57a_8 | m72_10 | m81_10 | m104_11 | m105a_11 | m110_11 | m131_11 | m183_11 | m185a_11 | |

**Table S7.** Populations from South America whose motifs were considered to elaborate the subset of motifs common to at least one of them and one African population (AmAfr subset).

| |
|---|
| Tarahumara,Warihio; Mai Huna (Coto); Chiriguano ; Opata; Cofan; Chacobo ; Yaqui,Mayo,Sinaloa; Napo,Kanelo; Ese'ejja ; Huichol; Waorani; Siriono ; Cora; Candoshi; Mojo,Manasi,etc.; Tepecano,Tepehuan; Zaparo; Parintintin; Durango Nahua; Shuar,Achuar,Huambiza; Tupari,Makurap,Ajuru; Tarascan; Aguaruna; Itene,Wari,other Rondonia; Aztec; Carijona; Surui,Zoro,Arua,Cinta Larga; Puebla & Huasteca Nahua; Barasana,Taibano,Makuna; Mundurucu; Otomi,Pame,Jonaz; Desana,Siriano,Tatuyo,Bara; Kamaiura; Huastec; Uanano,Tucano proper,Arapaso; Bakairi; Veracruz Nahuatl; Letuama,Ufaina,Yahuna; Kalapalo, Kuikuru; Popoluca; Cubeo; Waura,Mehinaku; Chinantec,Mazatec; Tariana; Trumai; Mixtec,Trique; Yucuna, Kabiyari; Rikbaktsa; Zapotec,Chatino; Macu; Kayabi; Oaxaca Mixe; Puinave; Iranxe; Chontal; Baniwa, Bare, Piapoco; Nambikwara; Zoque; Witoto,Ocaina; Paresi; Soconuzco Mestizos; Bora; Bororo; Tzotzil; Andoque; Umotina; Tseltal; Ticuna; Karaja; Chol; Yagua; Tapirape; Chorti; Omagua, Cocama; Cayapo; Yucatec, Itza; Chayahuita; Suya,Txukarramae; Lacandon; Urarina; Ramkokamekra,Apaniekra; Tzutujil; Katawishi,Manao etc.; Craho; Quiche,Pocomam; Mura; Crenye; Kanjobal,Chuj,Mocho; Mawe; Apinaye; Mam, Ixil; Juruna; Shavante; Pipil; Shipaya; Sherente; Lenca; Anambe; Kariri; Paya,Sumu,Misquito; Asurini,Paracana.; Ofaie; Rama,Guatuso; Lower Amazon; Guarani; Bribri,Cabecar; Tenetehara; Ache; Boruca; Urubu (Kaapor); Kaingang; Guaymi,Bocota; Tupinamba; Sheta; Kuna; Amuesha; Botocudo; Choco; Ashaninca; Kamakan; Kogi; Machiguenga; Maxakali; Ijka; Piro; Caduveo; Guajiro; Culina,Ipurina; Tereno; Chimila; Kanamari; Ayoreo; Bari; Shipibo-Conibo; Puelche; Tunebo; Cashibo; Tehuelche; Muisca; Marubo; Northern Peru: Sierra; Paez,Guambia; Cashinahua,Mayoruna; Lima dep.; Kamsa, Ingano; Harakmbet; Pasco,Junin,Huancavelica dep.; Cayapa (Chachi); Tacana; Ayacucho dep.; Colorado (Tsachila); Moseten,Chimane; Kechua: South Peru, Bolivia; Highland Ecuador; Yuracare; Aimara; Chipaya; Hoti; Akuriyo; Atacameno; Yanomamo; Waiwai; Yaruro; Sanema; Hixkaryana; Sicuani; Pemon; Kaxuyana; Cuiva; Akawai; Wayana, Aparai; Guayabero; Locono; Wayampi, Emerillon; Makiritare; Palikur; Siona,Coreguaje; Piaroa; Karina, Galibi; Chamacoco; Saliva; Orinoco Caribs; Nivakle; Yabarana; Makuxi; Chorote; Panare (E'napa); Wapishana; Mataco; Toba; Trio; Makka; Mocovi; Vilela; Lengua, Sanapana; Mapuche |

**Table S8**. Subsets of thematically linked motifs, with their source publication and the number of populations from our global dataset that included at least one of them.

| Subset | Motifs included | Global populations with >0 motifs |
|---|---|---|
| Cosmogonic motifs (d'Huy, 2018) | All motifs related to "The origins of the characteristics of the environment" in Berezkin's database (registered under the letter B). | All populations. |
| Motifs related to a Cosmogonic Dive hypothesised to have been present since the first diffusion of that myth (d'Huy, 2020b, pp.175-176) | b2a_3; b3a_3; b3b_3; b46c_2; b50_7; b77_3; b82_7; b87_2; c2_3; c3_7; c5a_3; c6_10; c6a_3; c6c_3; c6d_3; c19_3; c32_8 | Total: 66<br>Abkhasians, Adygei, Altaians, Armenians, Balkars, Bashkirs, Batak, Belarusians, Borneo_merged, Burmese, Buryats, Chukchis, Chuvash, Croats, Eskimo, Estonians, Evenkis_merged, Evens, Finns, French, Georgians, Germans, Gond, Gujaratis, Han, Hantis, Ho, Igorot, Ingrians, Iranians, Italians, Japanese, Karelian, Kazakhs, Komis, Koryaks_merged, Kyrgyzians, Latvians, Lezgins, Lithuanians, Malay, Mansis, Maris, Mayas, Mixtec, |

| | | |
|---|---|---|
| | | Mongolians_merged, Mordovians, Nenets_merged, Nganasans, North_Ossetians, Palestinians, Pamiris, Papuans, Poles, Portugese, Russians, Saudis, Shors, Tajiks, Teleut, Tuvinians, Udmurts, Ukranians, Vietnamese, Wichi, Yakuts |
| Motifs related to the Pleiades (d'Huy & Berezkin, 2017) | B42k; b47; b47a; b59; b60; i94; i95; i98a; i98b; i99; i100; i100a; i100c; i108; i114; i115a; i122; i130; m50 | Total: 43<br>Altaians, Armenians, Bashkirs, Belarusians, Chukchis, Chuvash, Croats, Eskimo, Estonians, Evens, Finns, French, Germans, Han, Hantis, Italians, Japanese, Kazakhs, Komis, Koryaks_merged, Kyrgyzians, Latvians, Lezgins, Lithuanians, Mansis, Maris, Mongolians_merged, Mordovians, Nenets_merged, Nganasans, North_Ossetians, Palestinians, Papuan, Poles, Russians, Saami_merged, Swedes, Tuvinians, Udmurts, Ukranians, Vepsas, Wichi, Yakuts |
| Motifs related to Matriarchy (d'Huy, 2017c) | F1; F5; F5A; F8; F16; F16B; F16C; F16D; F38; F40A; F40B; F41; F42; F42A; F43; F43A; F43B; F43C; F44; F45; F45A; F45B; F46; F46A; F47; F47A; F48; F97 | Total: 35<br>Abkhasians, Adygei, Altaians, Azeris, Balkars, Burmese, Chukchis, Chuvash, Eskimo, Finns, Georgians, Germans, Gond, Han, Hantis, Ho, Igorot, Iranians, Japanese, Kazakhs, Lezgins, Mansis, Maris, Mayas, Mongolians_merged, Nganasans, Palestinians, Papuans, Russians, Saami_merged, Saudis, Ukranians, Vietnamese, Wichi, Yakuts |
| Motifs related to After-death beliefs (d'Huy, 2020a) | a30_1; a36_4; b49_7; b71_3; d13h_8; d13hh_8; e14_6; e31a_11; e31b_11; f74_10; f94_10; h1a_4; h1b_4; h1bb_4; h1c_4; h2_4; h3_4; h4_4; h4a_4; h5_4; h6a_4; h6b_4; h6bb_4; h6c_4; h6d_4; h9b_4; h10_4; h11_8; h12_8; h12a_10; h12b_8; h13_10; h14_10; h16_8; h24b_4; h24c_4; h25_4; h31_4; h34a_4; h36_11; h36a_4; h36b_4; h36c_4; h36d_4; h36e_4; h36f_4; h36ff_4; h36g_4; h36gg_4; h36h_4; h36hh_4; h36I_4; h36j_4; h41_4; h42_4; h50_3; h52_10; h53_8; h56_4; i27_8; i27a_8; i30_8; i31_8; i33_8; i65_2; i96_3; i119_8 | Total: 62<br>Abkhasians, Adygei, Altaians, Armenians, Azeris, Balkars, Bashkirs, Belarusians, Borneo_merged, Burmese, Buryats, Chukchis, Chuvash, Eskimo, Estonians, Evenkis_merged, Evens, Finns, French, Georgians, Germans, Gond, Han, Hantis, Ho, Igorot, Iranians, Italians, Japanese, Kazakhs, Komis, Kyrgyzians, Latvians, Lithuanians, Malay, Mansis, Maris, Mayas, Mixtec, Mongolians_merged, Mordovians, Nenets_merged, Nganasans, North_Ossetians, Palestinians, Papuans, Poles, Portugese, Russians, Saami_merged, Saudis, Shors, Swedes, Tajiks, Teleut, Turkmens, Tuvinians, Udmurts, Ukranians, Vietnamese, Wichi, Yakuts |

**Mythological Admixture**

**Figure S1.** Cross-validation error values for different numbers (K) of components when running the software ADMIXTURE (Alexander et al., 2009).

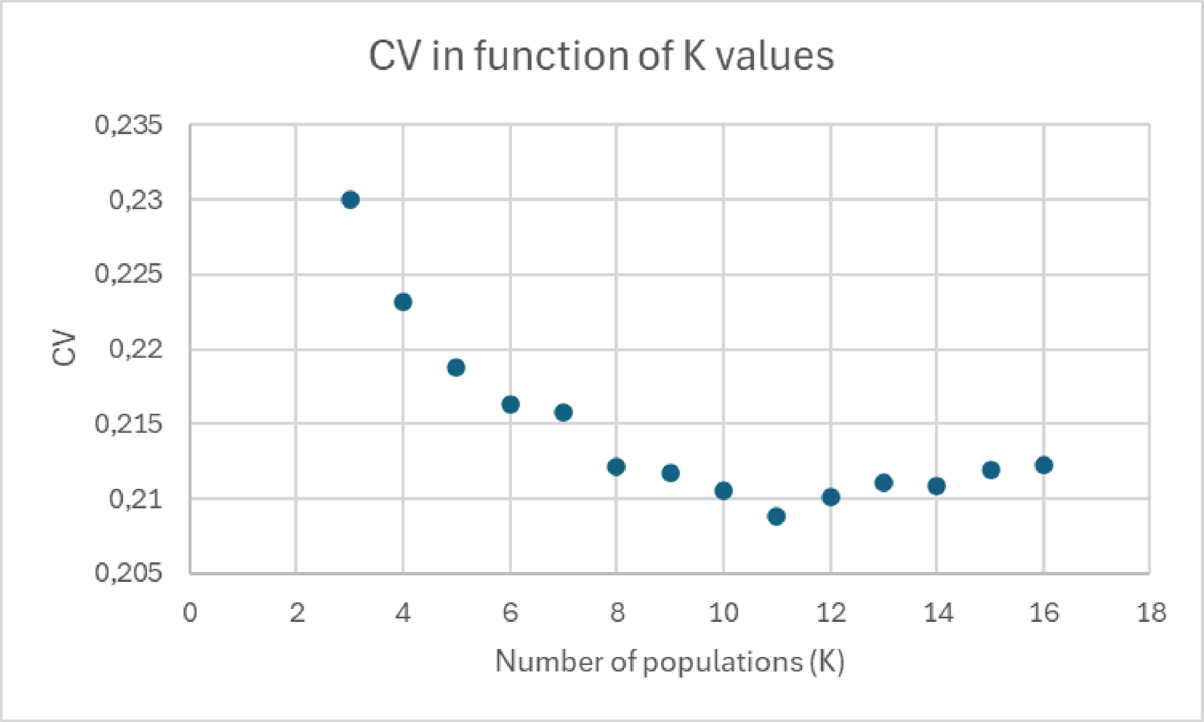


**Figure S2.** Admixture plot describing the composition of 781 mythological traditions around the world as the combination of 9 components.

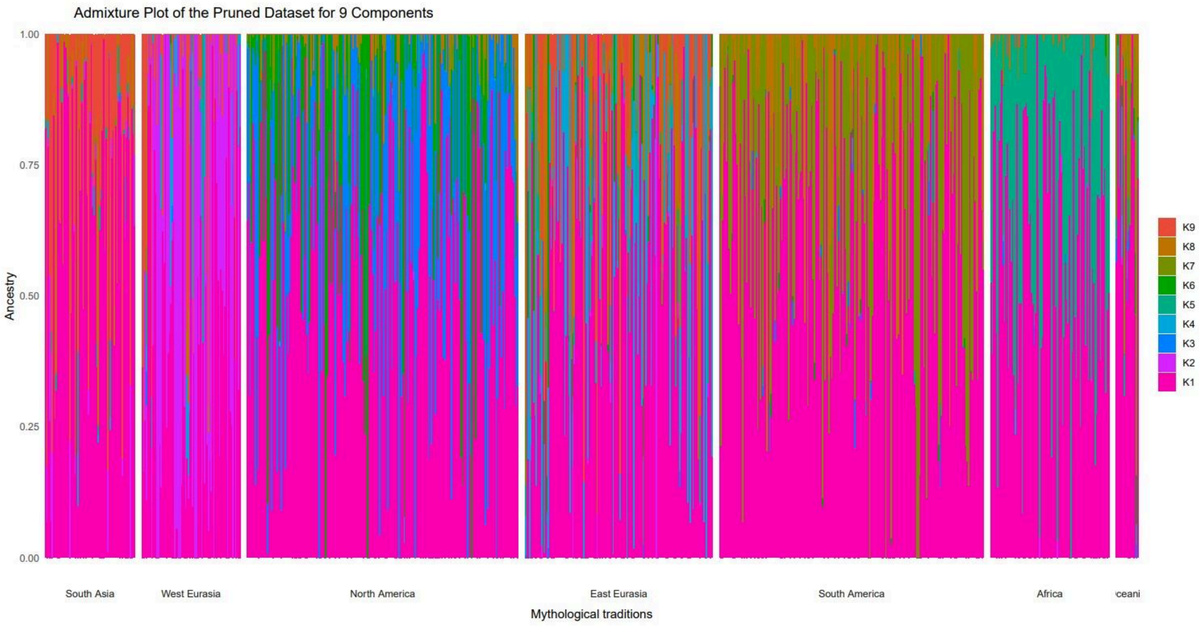

**Figure S3.** Barplot highlighting the presence of K1 in the 781 mythological traditions of our dataset.

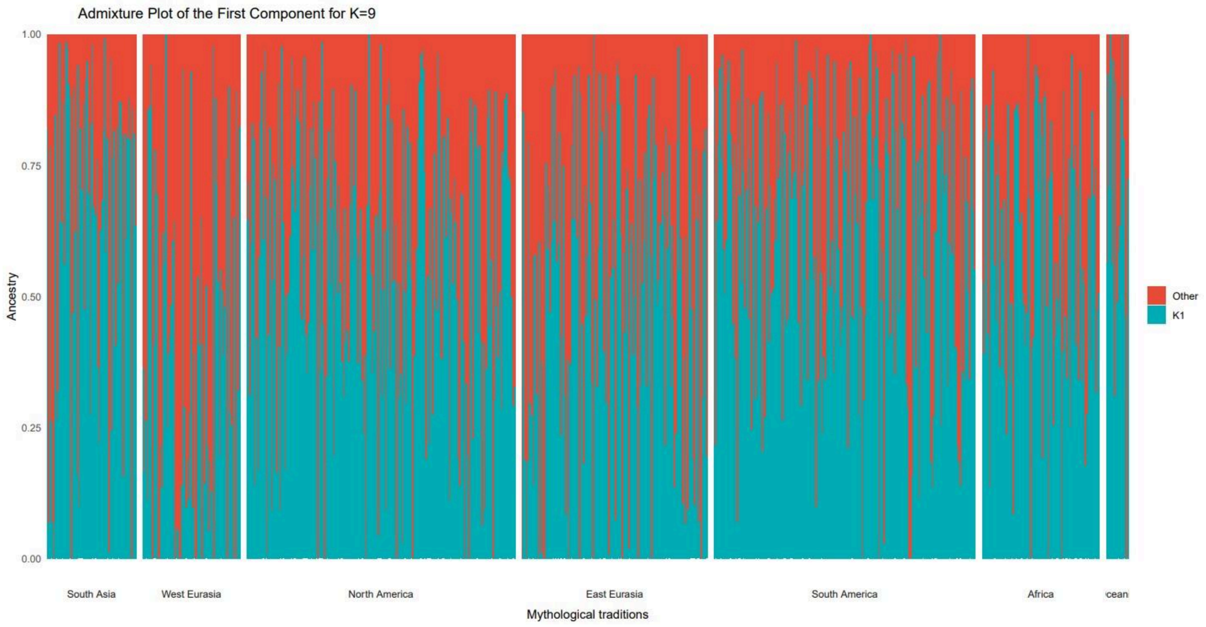


**Figure S4.** World map highlighting the distribution of K1 in the 781 mythological traditions present in our dataset.

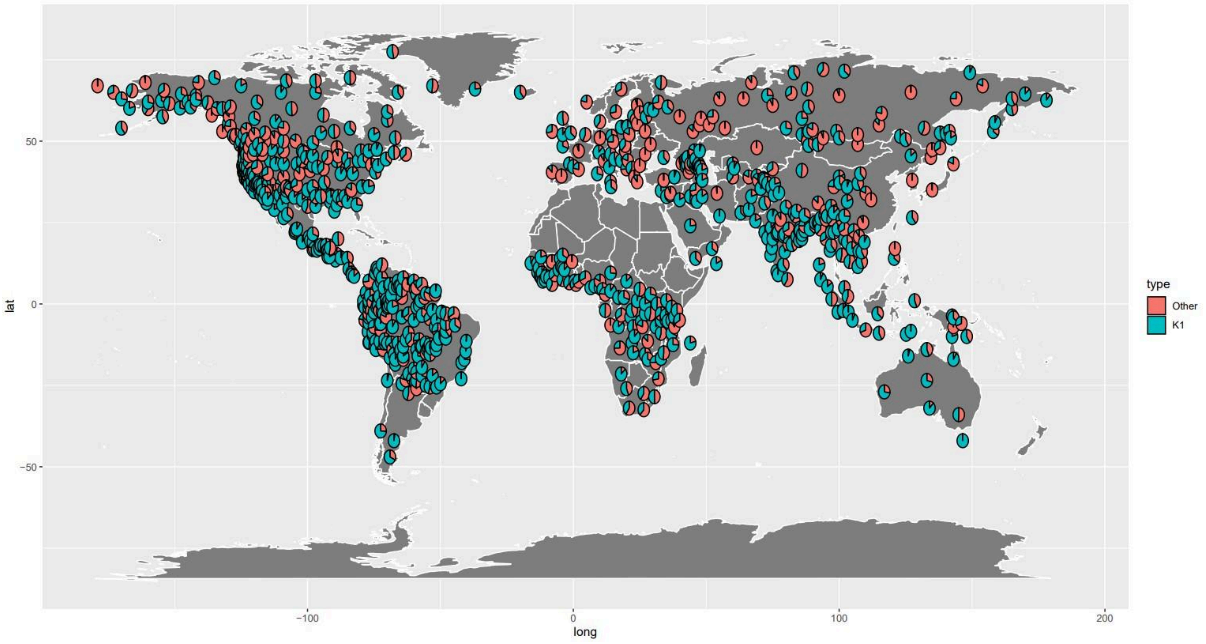


**Figure S5.** Barplot displaying ADMIXTURE results for the description of 781 worldwide mythological traditions according to 9 components, with K1 masked and K2-9 proportions rescaled to account for it in all mythological traditions, grouped per broad world region (SAs = South Asia, Weur = West Eurasia, NAm = North America, EEur = East Eurasia, SAm = South America, Afr = Africa, Oc = Oceania).

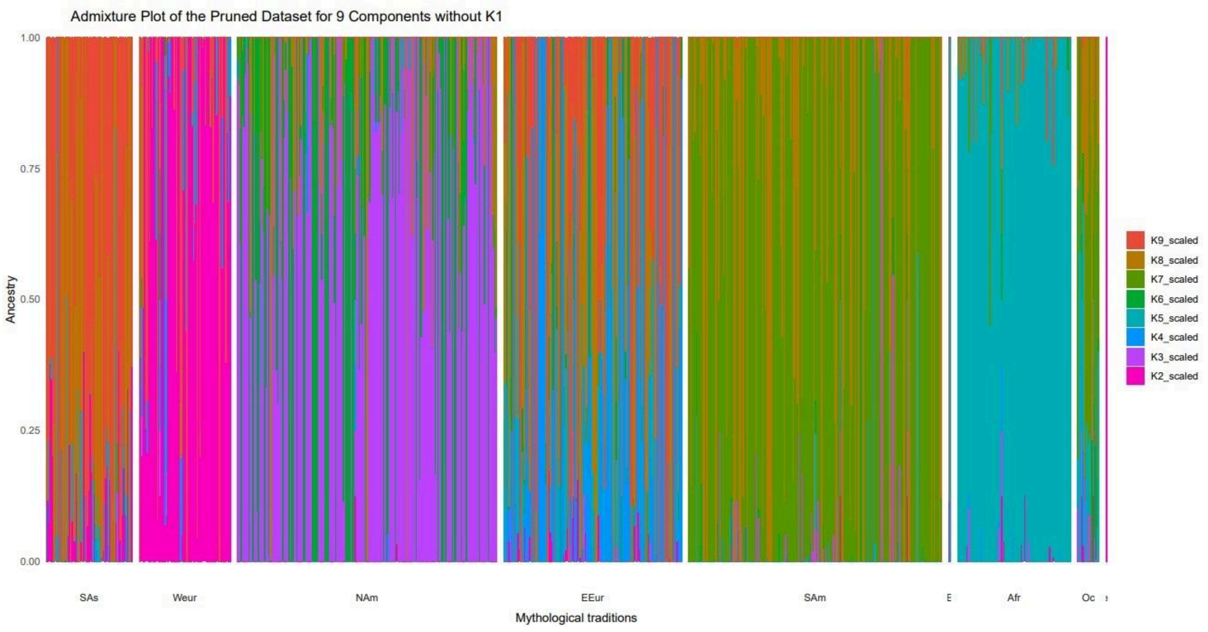


### Ethnolinguistic Structure

**Figure S6**. PCA of the 78 populations in our global dataset based on the Fst matrix.

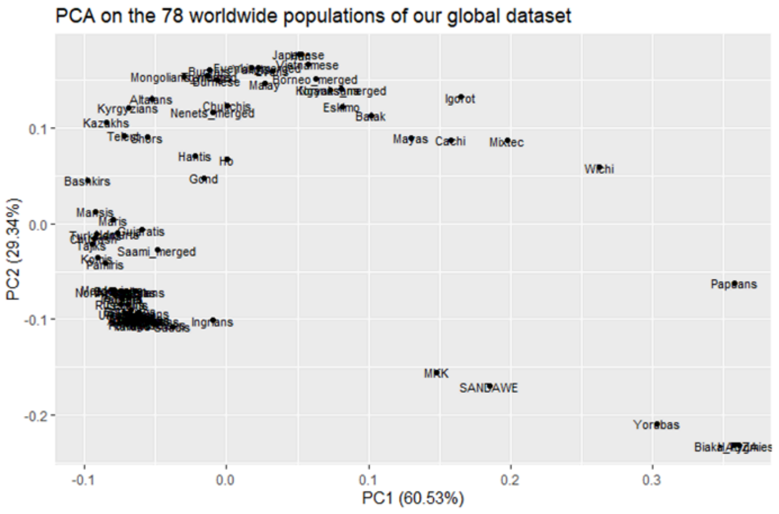

**Figure S7.** PCA on the 25 Eurasian populations we had in common with Bortolini et al. (2017)[1]

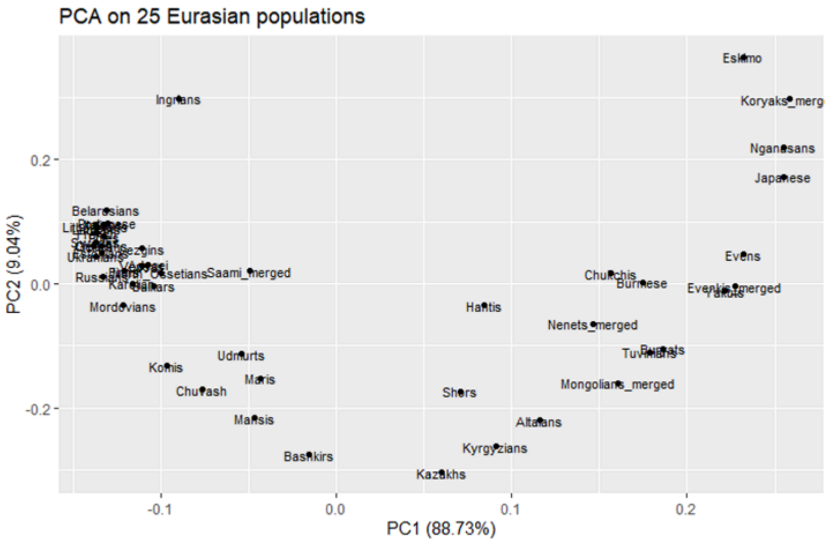


**Figure S8.** Boxplot representing the variability in the genetic distances between 25 Eurasian populations measured as with Fst (left) or average pairwise distance between two genomes coming from two different populations (right).

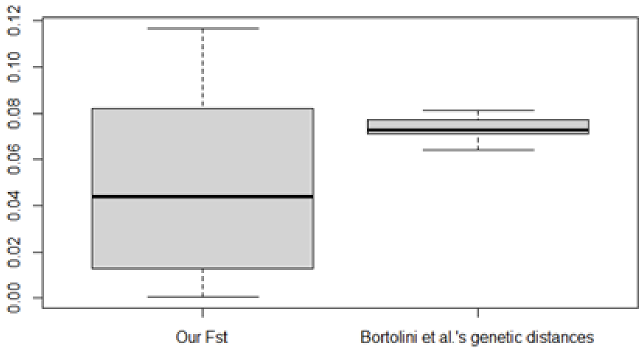


### Model comparison

**Figure S9.** Model comparison between demic diffusion (mythemic~genetic), cultural diffusion predicted by isolation-by-distance (mythemic~geographic), demic diffusion biassed by ethnolinguistic barriers (mythemicL~geneticL), cultural diffusion biassed by ethnolinguistic barriers (mythemicL~geographic) through the computation of Pearson's product moment correlation coefficient at different geographic bins (size = 2,000 km) over cumulative geographic distances.

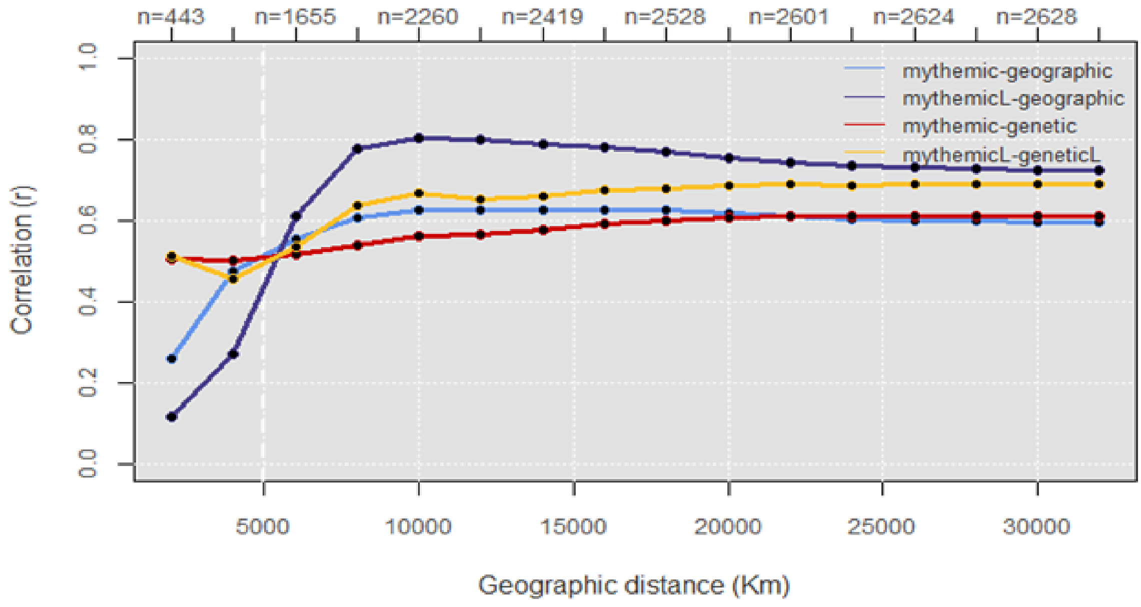


**Diffusion of thematically linked motifs**

In the studies we obtain our thematic groups from, Cosmogony- and Cosmogonic Dive-related motifs are hypothesised to have originated in Eurasia before spreading into the Americas during the Upper Palaeolithic [2–4]. On the other hand, imagery referring to the Pleiades, Matriarchy, and After-Death is conjectured to have been present in Africa and to have subsequently followed the first human migrations towards Eurasia [5–7].

Applying the methodology developed by Bortolini et al. (2017), we find that correlation values are inferior to those obtained for the whole dataset of motifs (Tables 1, 2, S9-S18) and always superior when we implement the correction for ethnolinguistic barriers. With Pearson's product-moment and bias-corrected correlations, both isolation by distance and past migrations appear to have played a role in shaping their current distribution worldwide, though the extent to which they do varies depending on the subset considered (Tables S9, S11, S13, S15, S17). Partial distance correlations, on the other hand, indicate a predominant role of cultural diffusion in the spread of motifs related to Cosmogony and to a Cosmogonic Dive (Tables S10 and S12). In the first case, a reduced phylogenetic signal had already been noted by d'Huy (2018) [4], suggesting a particularly cultural spread of that ensemble. In the second, when the impact of ethnolinguistic differentiation is neglected, partial distance correlations don't offer any statistical support for demic processes and don't either when it comes to motifs related to the Pleiades (Table SS.14). As for those having to do with Matriarchy, partial distance correlations are almost null and/or statistically insignificant (Table SS.16). The same is observed with the After-Death subset (Table SS.18), except when a linguistic correction is applied, in which case both models of cultural and demic diffusion are supported, with the first being prevalent.

**Table S9.** Model comparison between demic diffusion (mythemic~genetic), cultural diffusion predicted by isolation-by-distance (IBD) (mythemic~geographic), demic diffusion biased by ethnolinguistic barriers (mythemicL~geneticL), cultural diffusion biased by ethnolinguistic barriers (mythemicL~geographic). The values reported correspond to Pearson's product-moment correlations (cor), bias-corrected correlations (b.-c. cor), and their respective associated p-values. Mythological distances calculated were between non-African populations of the global dataset, using motifs of the Cosmogonic subset.

| Model | cor | p-value | b.-c. cor | p-value |
|---|---|---|---|---|
| Mythemic~genetic | 0.32 | < 0.001 | 0.32 | < 0.001 |
| Mythemic~geographic | 0.35 | < 0.001 | 0.4 | < 0.001 |
| MythemicL~geneticL | 0.6 | < 0.001 | 0.54 | < 0.001 |
| MythemicL~geographic | 0.66 | < 0.001 | 0.61 | < 0.001 |

**Table S10.** Partial distance correlations and their associated p-values for the different models under analysis. Mythological distances calculated were between non-African populations of the global dataset, using motifs of the Cosmogonic subset.

| Model | Partial correlation | p-value |
|---|---|---|
| Mythemic~genetic, geographic | 0.12 | < 0.001 |
| Mythemic~geographic, genetic | 0.16 | < 0.001 |
| MythemicL~geneticL, geographic | 0.2 | < 0.001 |
| MythemicL~geographic, geneticL | 0.38 | < 0.001 |

**Table S11.** Model comparison between demic diffusion (mythemic~genetic), cultural diffusion predicted by isolation-by-distance (IBD) (mythemic~geographic), demic diffusion biased by ethnolinguistic barriers (mythemicL~geneticL), cultural diffusion biased by ethnolinguistic barriers (mythemicL~geographic). The values reported correspond to Pearson's product-moment correlations (cor), bias-corrected correlations (b.-c. cor), and their respective associated p-values. Mythological distances calculated were between non-African populations of the global dataset, using motifs of the Cosmogonic Dive subset.

| Model | cor | p-value | b.-c. cor | p-value |
|---|---|---|---|---|
| Mythemic~genetic | 0.16 | < 0.001 | 0.12 | < 0.001 |
| Mythemic~geographic | 0.21 | < 0.001 | 0.15 | < 0.001 |
| MythemicL~geneticL | 0.38 | < 0.001 | 0.34 | < 0.001 |
| MythemicL~geographic | 0.43 | < 0.001 | 0.38 | < 0.001 |

**Table S12.** Partial distance correlations and their associated p-values for the different models under analysis. Mythological distances calculated were between non-African populations of the global dataset, using motifs of the Cosmogonic Dive subset.

| Model | Partial correlation | p-value |
|---|---|---|
| Mythemic~genetic, geographic | 0.01 | 0.26 |
| Mythemic~geographic, genetic | 0.09 | < 0.001 |
| MythemicL~geneticL, geographic | 0.11 | < 0.001 |
| MythemicL~geographic, geneticL | 0.22 | < 0.001 |

**Table S13.** Model comparison between demic diffusion (mythemic~genetic), cultural diffusion predicted by isolation-by-distance (IBD) (mythemic~geographic), demic diffusion biased by ethnolinguistic barriers (mythemicL~geneticL), cultural diffusion biased by ethnolinguistic barriers (mythemicL~geographic). The values reported correspond to Pearson's product-moment correlations (cor), bias-corrected correlations (b.-c. cor), and their respective associated p-values. Mythological distances calculated were between non-African populations of the global dataset, using motifs of the Pleiades subset.

| Model | cor | p-value | b.-c. cor | p-value |
|---|---|---|---|---|
| Mythemic~genetic | 0.06 | 0.0014 | 0.09 | < 0.001 |
| Mythemic~geographic | -0.03 | 0.1 | 0.16 | < 0.001 |
| MythemicL~geneticL | 0.18 | < 0.001 | 0.19 | < 0.001 |
| MythemicL~geographic | 0.16 | < 0.001 | 0.29 | < 0.001 |

**Table S14.** Partial distance correlations and their associated p-values for the different models under analysis. Mythological distances calculated were between non-African populations of the global dataset, using motifs of the Pleiades subset.

| Model | Partial correlation | p-value |
|---|---|---|
| Mythemic~genetic, geographic | -0.05 | 1 |
| Mythemic~geographic, genetic | 0.15 | < 0.001 |
| MythemicL~geneticL, geographic | -0.02 | 0.89 |
| MythemicL~geographic, geneticL | 0.23 | < 0.001 |

**Table S15.** Model comparison between demic diffusion (mythemic~genetic), cultural diffusion predicted by isolation-by-distance (IBD) (mythemic~geographic), demic diffusion biased by ethnolinguistic barriers (mythemicL~geneticL), cultural diffusion biased by ethnolinguistic barriers (mythemicL~geographic). The values reported correspond to Pearson's product-moment correlations (cor), bias-corrected correlations (b.-c. cor), and their respective associated p-values. Mythological distances calculated were between non-African populations of the global dataset, using motifs of the Matriarchy subset.

| Model | cor | p-value | b.-c. cor | p-value |
|---|---|---|---|---|
| Mythemic~genetic | 0.12 | < 0.001 | 0.01 | 0.27 |
| Mythemic~geographic | 0.07 | < 0.001 | 0.004 | 0.42 |
| MythemicL~geneticL | 0.21 | < 0.001 | 0.15 | < 0.001 |
| MythemicL~geographic | 0.18 | < 0.001 | 0.13 | < 0.001 |

**Table S16.** Partial distance correlations and their associated p-values for the different models under analysis. Mythological distances calculated were between non-African populations of the global dataset, using motifs of the Matriarchy subset.

| Model | Partial correlation | p-value |
|---|---|---|
| Mythemic~genetic, geographic | 0.01 | 0.18 |
| Mythemic~geographic, genetic | -0.007 | 0.58 |
| MythemicL~geneticL, geographic | 0.08 | < 0.001 |
| MythemicL~geographic, geneticL | 0.03 | 0.06 |

**Table S17.** Model comparison between demic diffusion (mythemic~genetic), cultural diffusion predicted by isolation-by-distance (IBD) (mythemic~geographic), demic diffusion biased by ethnolinguistic barriers (mythemicL~geneticL), cultural diffusion biased by ethnolinguistic barriers (mythemicL~geographic). The values reported correspond to Pearson's product-moment correlations (cor), bias-corrected correlations (b.-c. cor), and their respective associated p-values. Mythological distances calculated were between non-African populations of the global dataset, using motifs of the After-Death subset.

| Model | cor | p-value | b.-c. cor | p-value |
|---|---|---|---|---|
| Mythemic~genetic | 0.2 | < 0.001 | 0.16 | < 0.001 |
| Mythemic~geographic | 0.18 | < 0.001 | 0.16 | < 0.001 |
| MythemicL~geneticL | 0.4 | < 0.001 | 0.35 | < 0.001 |
| MythemicL~geographic | 0.42 | < 0.001 | 0.38 | < 0.001 |

**Table S18.** Partial distance correlations and their associated p-values for the different models under analysis. Mythological distances calculated were between non-African populations of the global dataset, using motifs of the After-Death subset.

| Model | Partial correlation | p-value |
|---|---|---|
| Mythemic~genetic, geographic | 0.07 | 0.007 |
| Mythemic~geographic, genetic | 0.06 | 0.01 |
| MythemicL~geneticL, geographic | 0.12 | < 0.001 |
| MythemicL~geographic, geneticL | 0.2 | < 0.001 |

**West Eurasia**

**Table S19.** Model comparison in West Eurasia between demic diffusion (mythemic~genetic), cultural diffusion predicted by isolation-by-distance (mythemic~geographic), demic diffusion biased by ethnolinguistic barriers (mythemicL~geneticL), cultural diffusion biased by ethnolinguistic barriers (mythemicL~geographic). The values reported correspond to Pearson's product-moment correlations (cor), bias-corrected correlations (b.-c. cor), and their respective associated p-values.

| Model | cor | p-value | b.-c. cor | p-value |
|---|---|---|---|---|
| Mythemic~genetic | 0.69 | < 0.001 | 0.51 | < 0.001 |
| Mythemic~geographic | 0.27 | < 0.001 | 0.53 | < 0.001 |
| Genetic~geographic | 0.25 | < 0.001 | 0.7 | < 0.001 |
| MythemicL~geneticL | 0.6 | < 0.001 | 0.56 | < 0.001 |
| MythemicL~geographic | 0.56 | < 0.001 | 0.51 | < 0.001 |
| GeneticL~geographic | 0.45 | < 0.001 | 0.49 | < 0.001 |

**Table S20.** Model comparison in West Eurasia. Partial distance correlations and their associated p-values for the different models under analysis: demic diffusion (mythemic~genetic), cultural diffusion predicted by isolation-by-distance (mythemic~geographic), demic diffusion biassed by ethnolinguistic barriers (mythemicL~geneticL), cultural diffusion biassed by ethnolinguistic barriers (mythemicL~geographic).

| Model | Partial correlation | p-value |
|---|---|---|
| Mythemic~genetic, geographic | 0.23 | 0.004 |
| Mythemic~geographic, genetic | 0.28 | 0.002 |
| MythemicL~geneticL, geographic | 0.41 | < 0.001 |
| MythemicL~geographic, geneticL | 0.32 | < 0.001 |

**Figure S10.** Model comparison in West Eurasia between demic diffusion (mythemic~genetic), cultural diffusion predicted by isolation-by-distance (IBD) (mythemic~geographic), demic diffusion biassed by ethnolinguistic barriers (mythemicL~geneticL), cultural diffusion biassed by ethnolinguistic barriers (mythemicL~geographic) through the computation of Pearson's product moment correlation coefficient at different geographic bins (size = 1,000 km) over cumulative geographic distances.

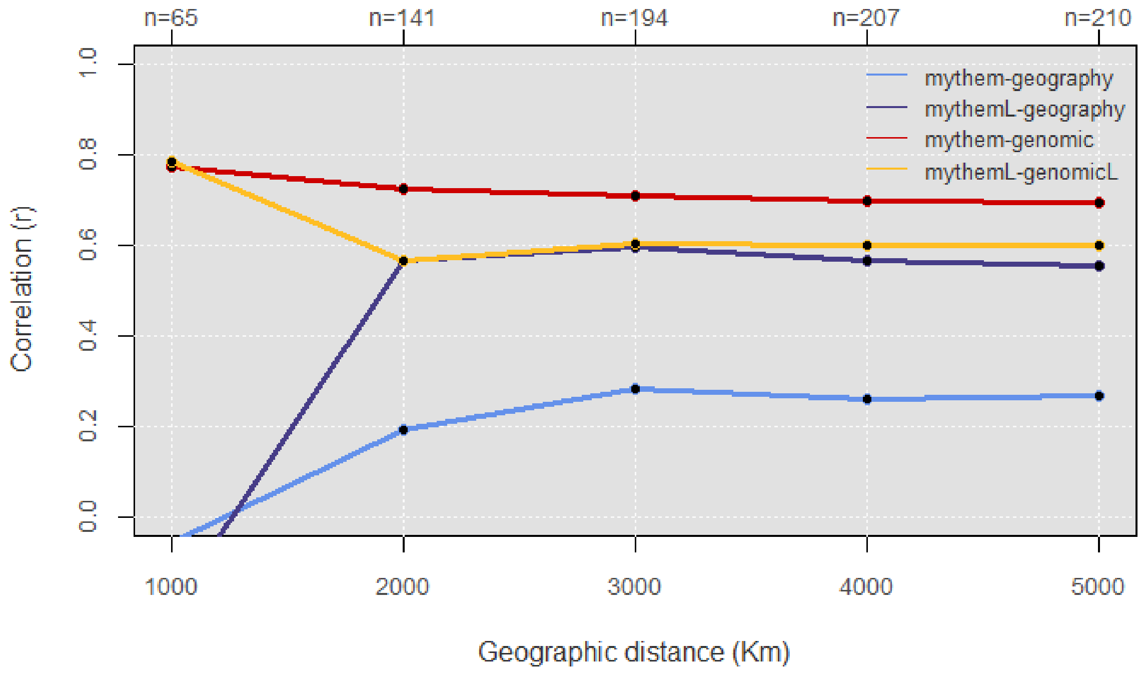

**Using subsets enriched in older motifs**

**Table S21.** Model comparison between demic diffusion (mythemic~genetic), cultural diffusion predicted by isolation-by-distance (IBD) (mythemic~geographic), demic diffusion biassed by ethnolinguistic barriers (mythemicL~geneticL), cultural diffusion biassed by ethnolinguistic barriers (mythemicL~geographic). The values reported correspond to Pearson's product-moment correlations (cor), bias-corrected correlations (b.-c. cor), and their respective associated p-values. Mythological distances were calculated between all non-African populations of the global dataset, using motifs present in at least one African and one South American populations.

| Model | cor | p-value | b.-c. cor | p-value |
|---|---|---|---|---|
| Mythemic~genetic | 0.48 | < 0.001 | 0.41 | < 0.001 |
| Mythemic~geographic | 0.56 | < 0.001 | 0.39 | < 0.001 |
| Genetic~geographic | 0.77 | < 0.001 | 0.73 | < 0.001 |
| MythemicL~geneticL | 0.65 | < 0.001 | 0.58 | < 0.001 |
| MythemicL~geographic | 0.72 | < 0.001 | 0.6 | < 0.001 |
| GeneticL~geographic | 0.84 | < 0.001 | 0.7 | < 0.001 |

**Table S22.** Partial distance correlations and their associated p-values for the different models under analysis. Mythological distances were calculated between all non-African populations of the global dataset, using motifs present in at least one African and one South American populations.

| Model | Partial correlation | p-value |
|---|---|---|
| Mythemic~genetic, geographic | 0.2 | < 0.001 |
| Mythemic~geographic, genetic | 0.15 | < 0.001 |
| MythemicL~geneticL, geographic | 0.27 | < 0.001 |
| MythemicL~geographic, geneticL | 0.34 | < 0.001 |

**Table S23.** Model comparison between demic diffusion (mythemic~genetic), cultural diffusion predicted by isolation-by-distance (IBD) (mythemic~geographic), demic diffusion biased by ethnolinguistic barriers (mythemicL~geneticL), cultural diffusion biased by ethnolinguistic barriers (mythemicL~geographic), using the OcAfr subset. The values reported correspond to Pearson's product-moment correlations (cor), bias-corrected correlations (b.-c. cor), and their respective associated p-values.

| Model | cor | p-value | b.-c. cor | p-value |
|---|---|---|---|---|
| Mythemic~genetic | 0.48 | < 0.001 | 0.39 | < 0.001 |
| Mythemic~geographic | 0.54 | < 0.001 | 0.37 | < 0.001 |
| Genetic~geographic | 0.77 | < 0.001 | 0.73 | < 0.001 |
| MythemicL~geneticL | 0.64 | < 0.001 | 0.57 | < 0.001 |
| MythemicL~geographic | 0.7 | < 0.001 | 0.59 | < 0.001 |
| GeneticL~geographic | 0.84 | < 0.001 | 0.7 | < 0.001 |

**Table S24.** Partial distance correlations and their associated p-values for the different models under analysis, using raw mythological distances between populations of the global dataset, computed for the OcAfr subset.

| Model | Partial correlation | p-value |
|---|---|---|
| Mythemic~genetic, geographic | 0.2 | < 0.001 |
| Mythemic~geographic, genetic | 0.13 | < 0.001 |
| MythemicL~geneticL, geographic | 0.28 | < 0.001 |
| MythemicL~geographic, geneticL | 0.32 | < 0.001 |

**Table S25.** Model comparison in Eurasia on the AmAfr subset between demic diffusion (mythemic~genetic), cultural diffusion predicted by isolation-by-distance (IBD) (mythemic~geographic), demic diffusion biased by ethnolinguistic barriers (mythemicL~geneticL), cultural diffusion biased by ethnolinguistic barriers (mythemicL~geographic). The values reported correspond to Pearson's product-moment correlations (cor), bias-corrected correlations (b.-c. cor), and their respective associated p-values.

| Model | cor | p-value | b.-c. cor | p-value |
|---|---|---|---|---|
| Mythemic~genetic | 0.45 | < 0.001 | 0.43 | < 0.001 |
| Mythemic~geographic | 0.46 | < 0.001 | 0.50 | < 0.001 |
| Genetic~geographic | 0.81 | < 0.001 | 0.86 | < 0.001 |

| | | | | |
|---|---|---|---|---|
| MythemicL~geneticL | 0.64 | < 0.001 | 0.65 | < 0.001 |
| MythemicL~geographic | 0.69 | < 0.001 | 0.65 | < 0.001 |
| GeneticL~geographic | 0.81 | < 0.001 | 0.72 | < 0.001 |

**Table S26.** Partial distance correlations and their associated p-values for the different models under analysis, using raw mythological distances between Eurasian populations, computed for the AmAfr subset.

| Model | Partial correlation | p-value |
|---|---|---|
| Mythemic~genetic, geographic | 0.024 | 0.16 |
| Mythemic~geographic, genetic | 0.27 | < 0.001 |
| MythemicL~geneticL, geographic | 0.35 | < 0.001 |
| MythemicL~geographic, geneticL | 0.34 | < 0.001 |

**Table S27.** Model comparison in Eurasia on the OcAfr subset between demic diffusion (mythemic~genetic), cultural diffusion predicted by isolation-by-distance (IBD) (mythemic~geographic), demic diffusion biased by ethnolinguistic barriers (mythemicL~geneticL), cultural diffusion biased by ethnolinguistic barriers (mythemicL~geographic). The values reported correspond to Pearson's product-moment correlations (cor), bias-corrected correlations (b.-c. cor), and their respective associated p-values.

| Model | cor | p-value | b.-c. cor | p-value |
|---|---|---|---|---|
| Mythemic~genetic | 0.46 | < 0.001 | 0.4 | < 0.001 |
| Mythemic~geographic | 0.45 | < 0.001 | 0.45 | < 0.001 |
| Genetic~geographic | 0.81 | < 0.001 | 0.86 | < 0.001 |
| MythemicL~geneticL | 0.63 | < 0.001 | 0.64 | < 0.001 |
| MythemicL~geographic | 0.68 | < 0.001 | 0.63 | < 0.001 |
| GeneticL~geographic | 0.81 | < 0.001 | 0.72 | < 0.001 |

**Table S28.** Partial distance correlations and their associated p-values for the different models under analysis, using raw mythological distances between Eurasian populations, computed for the OcAfr subset.

| Model | Partial correlation | p-value |
|---|---|---|
| Mythemic~genetic, geographic | 0.03 | 0.09 |
| Mythemic~geographic, genetic | 0.23 | < 0.001 |
| MythemicL~geneticL, geographic | 0.34 | < 0.001 |
| MythemicL~geographic, geneticL | 0.32 | < 0.001 |

**Time Groups**

**Table S29.** Model comparison between demic diffusion (mythemic~genetic) and cultural diffusion predicted by isolation-by-distance (IBD) (mythemic~geographic), using the pre-LGM dataset (Supplementary Data, 2.1). The values reported correspond to Pearson's product-moment correlations (cor), bias-corrected correlations (b.-c. cor), and their respective associated p-values.

| Model | cor | p-value | b.-c. cor | p-value |
|---|---|---|---|---|
| Mythemic~genetic | 0.71 | < 0.001 | 0.56 | < 0.001 |
| Mythemic~geographic | 0.81 | < 0.001 | 0.83 | < 0.001 |
| Genetic~geographic | 0.67 | < 0.001 | 0.42 | < 0.001 |

**Table S30.** Partial distance correlations and their associated p-values for the different models under analysis, using the pre-LGM dataset (Supplementary Data, 2.1).

| Model | Partial correlation | p-value |
|---|---|---|
| Mythemic~genetic, geographic | 0.41 | 0.002 |
| Mythemic~geographic, genetic | 0.78 | < 0.001 |

**Table S31.** Model comparison between demic diffusion (mythemic~genetic) and cultural diffusion predicted by isolation-by-distance (IBD) (mythemic~geographic), using the post-LGM/pre-Neolithic dataset (Supplementary Data, 2.2). The values reported correspond to Pearson's product-moment correlations (cor), bias-corrected correlations (b.-c. cor), and their respective associated p-values.

| Model | cor | p-value | b.-c. cor | p-value |
|---|---|---|---|---|
| Mythemic~genetic | 0.72 | < 0.001 | 0.41 | < 0.001 |
| Mythemic~geographic | 0.81 | < 0.001 | 0.83 | < 0.001 |
| Genetic~geographic | 0.72 | < 0.001 | 0.48 | < 0.001 |

**Table S32.** Partial distance correlations and their associated p-values for the different models under analysis, using the post-LGM/pre-Neolithic dataset (Supplementary Data, 2.2).

| Model | Partial correlation | p-value |
|---|---|---|
| Mythemic~genetic, geographic | 0.01 | 0.41 |
| Mythemic~geographic, genetic | 0.79 | < 0.001 |

**Table S33.** Model comparison between demic diffusion (mythemic~genetic) and cultural diffusion predicted by isolation-by-distance (IBD) (mythemic~geographic), using the Neolithic dataset (Supplementary Data, 2.3). The values reported correspond to Pearson's product-moment correlations (cor), bias-corrected correlations (b.-c. cor), and their respective associated p-values.

| Model | cor | p-value | b.-c. cor | p-value |
|---|---|---|---|---|
| Mythemic~genetic | 0.56 | < 0.001 | 0.32 | 0.002 |
| Mythemic~geographic | 0.81 | < 0.001 | 0.83 | < 0.001 |
| Genetic~geographic | 0.71 | < 0.001 | 0.46 | < 0.001 |

**Table S34.** Partial distance correlations and their associated p-values for the different models under analysis, using the Neolithic dataset (Supplementary Data, 2.3).

| Model | Partial correlation | p-value |
|---|---|---|
| Mythemic~genetic, geographic | -0.13 | 0.88 |
| Mythemic~geographic, genetic | 0.81 | < 0.001 |

**Table S35.** Model comparison between demic diffusion (mythemic~genetic) and cultural diffusion predicted by isolation-by-distance (IBD) (mythemic~geographic), using the Bronze Age dataset (Supplementary Data, 2.4). The values reported correspond to Pearson's product-moment correlations (cor), bias-corrected correlations (b.-c. cor), and their respective associated p-values.

| Model | cor | p-value | b.-c. cor | p-value |
|---|---|---|---|---|
| Mythemic~genetic | 0.72 | < 0.001 | 0.24 | 0.017 |
| Mythemic~geographic | 0.81 | < 0.001 | 0.83 | < 0.001 |
| Genetic~geographic | 0.76 | < 0.001 | 0.3 | 0.004 |

**Table S36.** Partial distance correlations and their associated p-values for the different models under analysis, using the Bronze Age dataset (Supplementary Data, 2.4).

| Model | Partial correlation | p-value |
|---|---|---|
| Mythemic~genetic, geographic | -0.01 | 0.47 |
| Mythemic~geographic, genetic | 0.81 | < 0.001 |

**Table S37.** Model comparison between demic diffusion (mythemic~genetic) and cultural diffusion predicted by isolation-by-distance (IBD) (mythemic~geographic), using the Iron Age dataset (Supplementary Data, 2.5). The values reported correspond to Pearson's product-moment correlations (cor), bias-corrected correlations (b.-c. cor), and their respective associated p-values.

| Model | cor | p-value | b.-c. cor | p-value |
|---|---|---|---|---|
| Mythemic~genetic | 0.59 | < 0.001 | 0.36 | < 0.001 |
| Mythemic~geographic | 0.81 | < 0.001 | 0.83 | < 0.001 |
| Genetic~geographic | 0.74 | < 0.001 | 0.48 | < 0.001 |

**Table S38.** Partial distance correlations and their associated p-values for the different models under analysis, using the Iron Age dataset (Supplementary Data, 2.5).

| Model | Partial correlation | p-value |
|---|---|---|
| Mythemic~genetic, geographic | -0.08 | 0.75 |

| | | |
|---|---|---|
| Mythemic~geographic, genetic | 0.8 | < 0.001 |

**Table S39.** Model comparison between demic diffusion (mythemic~genetic) and cultural diffusion predicted by isolation-by-distance (IBD) (mythemic~geographic), using 14 present-day populations (Supplementary Data, 2.6). The values reported correspond to Pearson's product-moment correlations (cor), bias-corrected correlations (b.-c. cor), and their respective associated p-values.

| Model | cor | p-value | b.-c. cor | p-value |
|---|---|---|---|---|
| Mythemic~genetic | 0.74 | < 0.001 | 0.71 | < 0.001 |
| Mythemic~geographic | 0.81 | < 0.001 | 0.83 | < 0.001 |
| Genetic~geographic | 0.86 | < 0.001 | 0.86 | < 0.001 |

**Table S40.** Partial distance correlations and their associated p-values for the different models under analysis, using 14 present-day populations (Supplementary Data, 2.6).

| Model | Partial correlation | p-value |
|---|---|---|
| Mythemic~genetic, geographic | 0.008 | 0.51 |
| Mythemic~geographic, genetic | 0.6 | < 0.001 |

**Table S41.** Model comparison between demic diffusion (mythemic~genetic) and cultural diffusion predicted by isolation-by-distance (IBD) (mythemic~geographic), using the pre-LGM dataset (Supplementary Data, 2.1) and AmAfr motifs (Table SS5). The values reported correspond to Pearson's product-moment correlations (cor), bias-corrected correlations (b.-c. cor), and their respective associated p-values.

| Model | cor | p-value | b.-c. cor | p-value |
|---|---|---|---|---|
| Mythemic~genetic | 0.31 | 0.002 | 0.48 | < 0.001 |
| Mythemic~geographic | 0.39 | < 0.001 | 0.45 | < 0.001 |
| Genetic~geographic | 0.67 | < 0.001 | 0.42 | < 0.001 |

**Table S42.** Partial distance correlations and their associated p-values for the different models under analysis, using the pre-LGM dataset (Supplementary Data, 2.1) and AmAfr motifs (Table SS5).

| Model | Partial correlation | p-value |
|---|---|---|

| | | |
|---|---|---|
| Mythemic~genetic, geographic | 0.36 | < 0.001 |
| Mythemic~geographic, genetic | 0.31 | 0.003 |

**Table S43.** Model comparison between demic diffusion (mythemic~genetic) and cultural diffusion predicted by isolation-by-distance (IBD) (mythemic~geographic), using the pre-LGM dataset (Supplementary Data, 2.1) and OcAfr motifs (Table SS6). The values reported correspond to Pearson's product-moment correlations (cor), bias-corrected correlations (b.-c. cor), and their respective associated p-values.

| Model | cor | p-value | b.-c. cor | p-value |
|---|---|---|---|---|
| Mythemic~genetic | 0.53 | 0.002 | 0.43 | < 0.001 |
| Mythemic~geographic | 0.61 | < 0.001 | 0.48 | < 0.001 |
| Genetic~geographic | 0.67 | < 0.001 | 0.42 | < 0.001 |

**Table S44.** Partial distance correlations and their associated p-values for the different models under analysis, using the pre-LGM dataset (Supplementary Data, 2.1) and OcAfr motifs (Table SS6).

| Model | Partial correlation | p-value |
|---|---|---|
| Mythemic~genetic, geographic | 0.28 | 0.006 |
| Mythemic~geographic, genetic | 0.36 | 0.003 |

**Table S45.** Model comparison between demic diffusion (mythemic~genetic) and cultural diffusion predicted by isolation-by-distance (IBD) (mythemic~geographic), using the post-LGM/pre-Neolithic dataset (Supplementary Data, 2.2) and AmAfr motifs (Table SS5). The values reported correspond to Pearson's product-moment correlations (cor), bias-corrected correlations (b.-c. cor), and their respective associated p-values.

| Model | cor | p-value | b.-c. cor | p-value |
|---|---|---|---|---|
| Mythemic~genetic | 0.45 | 0.002 | 0.14 | 0.11 |
| Mythemic~geographic | 0.39 | < 0.001 | 0.45 | < 0.001 |
| Genetic~geographic | 0.72 | < 0.001 | 0.48 | < 0.001 |

**Table S46.** Partial distance correlations and their associated p-values for the different models under analysis, using the post-LGM/pre-Neolithic dataset (Supplementary Data, 2.2) and AmAfr motifs (Table SS5).

| Model | Partial correlation | p-value |
|---|---|---|
| Mythemic~genetic, geographic | -0.1 | 0.81 |
| Mythemic~geographic, genetic | 0.44 | <0.001 |

**Table S47.** Model comparison between demic diffusion (mythemic~genetic) and cultural diffusion predicted by isolation-by-distance (IBD) (mythemic~geographic), using the Neolithic dataset (Supplementary Data, 2.3) and AmAfr motifs (Table SS5). The values reported correspond to Pearson's product-moment correlations (cor), bias-corrected correlations (b.-c. cor), and their respective associated p-values.

| Model | cor | p-value | b.-c. cor | p-value |
|---|---|---|---|---|
| Mythemic~genetic | 0.3 | 0.004 | 0.2 | 0.037 |
| Mythemic~geographic | 0.39 | < 0.001 | 0.45 | < 0.001 |
| Genetic~geographic | 0.71 | < 0.001 | 0.46 | < 0.001 |

**Table S48.** Partial distance correlations and their associated p-values for the different models under analysis, using the Neolithic dataset (Supplementary Data, 2.3) and AmAfr motifs (Table SS5).

| Model | Partial correlation | p-value |
|---|---|---|
| Mythemic~genetic, geographic | -0.005 | 0.47 |
| Mythemic~geographic, genetic | 0.41 | <0.001 |

**Table S49.** Model comparison between demic diffusion (mythemic~genetic) and cultural diffusion predicted by isolation-by-distance (IBD) (mythemic~geographic), using the Bronze Age dataset (Supplementary Data, 2.4) and AmAfr motifs (Table SS5). The values reported correspond to Pearson's product-moment correlations (cor), bias-corrected correlations (b.-c. cor), and their respective associated p-values.

| Model | cor | p-value | b.-c. cor | p-value |
|---|---|---|---|---|
| Mythemic~genetic | 0.34 | < 0.001 | 0.08 | 0.25 |
| Mythemic~geographic | 0.39 | < 0.001 | 0.45 | < 0.001 |
| Genetic~geographic | 0.76 | < 0.001 | 0.3 | 0.005 |

**Table S50.** Partial distance correlations and their associated p-values for the different models under analysis, using the Bronze Age dataset (Supplementary Data, 2.4) and AmAfr motifs (Table SS5).

| Model | Partial correlation | p-value |
|---|---|---|
| Mythemic~genetic, geographic | -0.06 | 0.72 |
| Mythemic~geographic, genetic | 0.45 | <0.001 |

**Table S51.** Model comparison between demic diffusion (mythemic~genetic) and cultural diffusion predicted by isolation-by-distance (IBD) (mythemic~geographic), using the Iron Age dataset (Supplementary Data, 2.5) and AmAfr motifs (Table SS5). The values reported correspond to Pearson's product-moment correlations (cor), bias-corrected correlations (b.-c. cor), and their respective associated p-values.

| Model | cor | p-value | b.-c. cor | p-value |
|---|---|---|---|---|
| Mythemic~genetic | 0.31 | 0.003 | 0.24 | 0.02 |
| Mythemic~geographic | 0.39 | < 0.001 | 0.45 | < 0.001 |
| Genetic~geographic | 0.74 | < 0.001 | 0.48 | < 0.001 |

**Table S52.** Partial distance correlations and their associated p-values for the different models under analysis, using the Iron Age dataset (Supplementary Data, 2.5) and AmAfr motifs (Table SS5).

| Model | Partial correlation | p-value |
|---|---|---|
| Mythemic~genetic, geographic | 0.03 | 0.36 |
| Mythemic~geographic, genetic | 0.39 | <0.001 |

**Table S53.** Model comparison between demic diffusion (mythemic~genetic) and cultural diffusion predicted by isolation-by-distance (IBD) (mythemic~geographic), using the present-day dataset (Supplementary Data, 2.6) and AmAfr motifs (Table SS5). The values reported correspond to Pearson's product-moment correlations (cor), bias-corrected correlations (b.-c. cor), and their respective associated p-values.

| Model | cor | p-value | b.-c. cor | p-value |
|---|---|---|---|---|
| Mythemic~genetic | 0.43 | < 0.001 | 0.41 | < 0.001 |
| Mythemic~geographic | 0.39 | < 0.001 | 0.45 | < 0.001 |
| Genetic~geographic | 0.86 | < 0.001 | 0.86 | < 0.001 |

**Table S54.** Partial distance correlations and their associated p-values for the different models under analysis, using the present-day dataset (Supplementary Data, 2.6) and AmAfr motifs (Table SS5).

| Model | Partial correlation | p-value |
|---|---|---|
| Mythemic~genetic, geographic | 0.05 | 0.33 |
| Mythemic~geographic, genetic | 0.21 | 0.03 |

**Table S55.** Model comparison between demic diffusion (mythemic~genetic) and cultural diffusion predicted by isolation-by-distance (IBD) (mythemic~geographic), using only non-European traditions, except for "Germans", in the pre-LGM dataset (Supplementary Data, 2.1). The values reported correspond to Pearson's product-moment correlations (cor), bias-corrected correlations (b.-c. cor), and their respective associated p-values.

| Model | cor | p-value | b.-c. cor | p-value |
|---|---|---|---|---|
| Mythemic~genetic | 0.45 | 0.006 | 0.2 | 0.15 |
| Mythemic~geographic | 0.56 | < 0.001 | 0.66 | < 0.001 |
| Genetic~geographic | 0.38 | 0.02 | 0.23 | 0.12 |

**Table S56.** Partial distance correlations and their associated p-values for the different models under analysis, using only non-European traditions, except for "Germans", the pre-LGM dataset (Supplementary Data, 2.1).

| Model | Partial correlation | p-value |
|---|---|---|
| Mythemic~genetic, geographic | 0.06 | 0.34 |
| Mythemic~geographic, genetic | 0.64 | 0.003 |

**Table S57.** Model comparison between demic diffusion (mythemic~genetic) and cultural diffusion predicted by isolation-by-distance (IBD) (mythemic~geographic), using only non-European traditions, except for "Germans", in the post-LGM/pre-Neolithic dataset (Supplementary Data, 2.2). The values reported correspond to Pearson's product-moment correlations (cor), bias-corrected correlations (b.-c. cor), and their respective associated p-values.

| Model | cor | p-value | b.-c. cor | p-value |
|---|---|---|---|---|
| Mythemic~genetic | 0.29 | 0.08 | 0.6 | < 0.001 |
| Mythemic~geographic | 0.56 | < 0.001 | 0.66 | < 0.001 |
| Genetic~geographic | 0.55 | < 0.001 | 0.41 | 0.02 |

**Table S58.** Partial distance correlations and their associated p-values for the different models under analysis, using only non-European traditions, except for "Germans", in the post-LGM/pre-Neolithic dataset (Supplementary Data, 2.2).

| Model | Partial correlation | p-value |
|---|---|---|
| Mythemic~genetic, geographic | 0.48 | 0.003 |
| Mythemic~geographic, genetic | 0.57 | < 0.001 |

**Table S59.** Model comparison between demic diffusion (mythemic~genetic) and cultural diffusion predicted by isolation-by-distance (IBD) (mythemic~geographic), using only non-European traditions, except for "Germans", in the Neolithic dataset (Supplementary Data, 2.3). The values reported correspond to Pearson's product-moment correlations (cor), bias-corrected correlations (b.-c. cor), and their respective associated p-values.

| Model | cor | p-value | b.-c. cor | p-value |
|---|---|---|---|---|
| Mythemic~genetic | 0.11 | 0.51 | 0.1 | 0.31 |
| Mythemic~geographic | 0.56 | < 0.001 | 0.66 | < 0.001 |
| Genetic~geographic | 0.54 | < 0.001 | 0.25 | 0.09 |

**Table S60.** Partial distance correlations and their associated p-values for the different models under analysis, using only non-European traditions, except for "Germans", in the Neolithic dataset (Supplementary Data, 2.3).

| Model | Partial correlation | p-value |
|---|---|---|
| Mythemic~genetic, geographic | -0.09 | 0.69 |
| Mythemic~geographic, genetic | 0.66 | < 0.001 |

**Table S61.** Model comparison between demic diffusion (mythemic~genetic) and cultural diffusion predicted by isolation-by-distance (IBD) (mythemic~geographic), using only non-European traditions, except for "Germans", in the Bronze Age dataset (Supplementary Data, 2.4). The values reported correspond to Pearson's product-moment correlations (cor), bias-corrected correlations (b.-c. cor), and their respective associated p-values.

| Model | cor | p-value | b.-c. cor | p-value |
|---|---|---|---|---|
| Mythemic~genetic | 0.28 | 0.1 | 0.004 | 0.49 |
| Mythemic~geographic | 0.56 | < 0.001 | 0.66 | < 0.001 |
| Genetic~geographic | 0.52 | 0.001 | 0.1 | 0.31 |

**Table S62.** Partial distance correlations and their associated p-values for the different models under analysis, using only non-European traditions, except for "Germans", in the Bronze Age dataset (Supplementary Data, 2.4).

| Model | Partial correlation | p-value |
|---|---|---|
| Mythemic~genetic, geographic | -0.08 | 0.69 |
| Mythemic~geographic, genetic | 0.66 | 0.003 |

**Table S63.** Model comparison between demic diffusion (mythemic~genetic) and cultural diffusion predicted by isolation-by-distance (IBD) (mythemic~geographic), using only non-European traditions, except for "Germans", in the Iron Age dataset (Supplementary Data, 2.5). The values reported correspond to Pearson's product-moment correlations (cor), bias-corrected correlations (b.-c. cor), and their respective associated p-values.

| Model | cor | p-value | b.-c. cor | p-value |
|---|---|---|---|---|
| Mythemic~genetic | 0.16 | 0.34 | 0.14 | 0.23 |
| Mythemic~geographic | 0.56 | < 0.001 | 0.66 | < 0.001 |
| Genetic~geographic | 0.54 | < 0.001 | 0.29 | 0.07 |

**Table S64.** Partial distance correlations and their associated p-values for the different models under analysis, using only non-European traditions, except for "Germans", in the Iron Age dataset (Supplementary Data, 2.5).

| Model | Partial correlation | p-value |
|---|---|---|
| Mythemic~genetic, geographic | -0.07 | 0.64 |
| Mythemic~geographic, genetic | 0.65 | 0.003 |

**Table S65.** Model comparison between demic diffusion (mythemic~genetic) and cultural diffusion predicted by isolation-by-distance (IBD) (mythemic~geographic), using only non-European traditions, except for "Germans", in the 14 present-day populations (Supplementary Data, 2.6). The values reported correspond to Pearson's product-moment correlations (cor), bias-corrected correlations (b.-c. cor), and their respective associated p-values.

| Model | cor | p-value | b.-c. cor | p-value |
|---|---|---|---|---|
| Mythemic~genetic | 0.58 | < 0.001 | 0.54 | 0.002 |
| Mythemic~geographic | 0.56 | < 0.001 | 0.66 | < 0.001 |
| Genetic~geographic | 0.68 | < 0.001 | 0.74 | < 0.001 |

**Table S66.** Partial distance correlations and their associated p-values for the different models under analysis, using only non-European traditions, except for "Germans", in the 14 present-day populations (Supplementary Data, 2.6).

| Model | Partial correlation | p-value |
|---|---|---|
| Mythemic~genetic, geographic | 0.1 | 0.33 |
| Mythemic~geographic, genetic | 0.46 | 0.005 |

**Table S67.** Model comparison between demic diffusion (mythemic~genetic) and cultural diffusion predicted by isolation-by-distance (IBD) (mythemic~geographic), using only non-European traditions, except for "Germans", in the pre-LGM dataset (Supplementary Data, 2.1) and AmAfr motifs (Table SS6). The values reported correspond to Pearson's product-moment correlations (cor), bias-corrected correlations (b.-c. cor), and their respective associated p-values.

| Model | cor | p-value | b.-c. cor | p-value |
|---|---|---|---|---|
| Mythemic~genetic | 0.47 | 0.004 | 0.42 | 0.01 |
| Mythemic~geographic | 0.26 | 0.13 | 0.47 | 0.006 |
| Genetic~geographic | 0.38 | 0.02 | 0.23 | 0.12 |

**Table S68.** Partial distance correlations and their associated p-values for the different models under analysis, using only non-European traditions, except for "Germans", in the pre-LGM dataset (Supplementary Data, 2.1) and AmAfr motifs (Table SS6).

| Model | Partial correlation | p-value |
|---|---|---|
| Mythemic~genetic, geographic | 0.36 | 0.05 |
| Mythemic~geographic, genetic | 0.42 | 0.008 |

**Table S69.** Model comparison between demic diffusion (mythemic~genetic) and cultural diffusion predicted by isolation-by-distance (IBD) (mythemic~geographic), using only non-European traditions, except for "Germans", in the post-LGM/pre-Neolithic dataset (Supplementary Data, 2.2) and AmAfr motifs (Table SS5). The values reported correspond to Pearson's product-moment correlations (cor), bias-corrected correlations (b.-c. cor), and their respective associated p-values.

| Model | cor | p-value | b.-c. cor | p-value |
|---|---|---|---|---|
| Mythemic~genetic | 0.38 | 0.02 | 0.43 | 0.01 |
| Mythemic~geographic | 0.26 | 0.13 | 0.47 | 0.006 |
| Genetic~geographic | 0.55 | < 0.001 | 0.41 | 0.02 |

**Table S70.** Partial distance correlations and their associated p-values for the different models under analysis, using only non-European traditions, except for "Germans", in the post-LGM/pre-Neolithic dataset (Supplementary Data, 2.2) and AmAfr motifs (Table SS5).

| Model | Partial correlation | p-value |
|---|---|---|
| Mythemic~genetic, geographic | 0.29 | 0.06 |
| Mythemic~geographic, genetic | 0.26 | 0.03 |

**Table S71.** Model comparison between demic diffusion (mythemic~genetic) and cultural diffusion predicted by isolation-by-distance (IBD) (mythemic~geographic), using only non-European traditions, except for "Germans", in the Neolithic dataset (Supplementary Data, 2.3) and AmAfr motifs (Table SS5). The values reported correspond to Pearson's product-moment correlations (cor), bias-corrected correlations (b.-c. cor), and their respective associated p-values.

| Model | cor | p-value | b.-c. cor | p-value |
|---|---|---|---|---|
| Mythemic~genetic | 0.21 | 0.21 | 0.16 | 0.21 |
| Mythemic~geographic | 0.26 | 0.13 | 0.47 | 0.006 |
| Genetic~geographic | 0.54 | < 0.001 | 0.25 | 0.1 |

**Table S72.** Partial distance correlations and their associated p-values for the different models under analysis, using only non-European traditions, except for "Germans", in the Neolithic dataset (Supplementary Data, 2.3) and AmAfr motifs (Table SS5).

| Model | Partial correlation | p-value |
|---|---|---|
| Mythemic~genetic, geographic | 0.04 | 0.34 |
| Mythemic~geographic, genetic | 0.45 | 0.007 |

**Table S73.** Model comparison between demic diffusion (mythemic~genetic) and cultural diffusion predicted by isolation-by-distance (IBD) (mythemic~geographic), using only non-European traditions, except for "Germans", in the Bronze Age dataset (Supplementary Data, 2.4) and AmAfr motifs (Table SS5). The values reported correspond to Pearson's product-moment correlations (cor), bias-corrected correlations (b.-c. cor), and their respective associated p-values.

| Model | cor | p-value | b.-c. cor | p-value |
|---|---|---|---|---|
| Mythemic~genetic | 0.22 | 0.2 | -0.009 | 0.52 |
| Mythemic~geographic | 0.26 | 0.13 | 0.47 | 0.006 |
| Genetic~geographic | 0.52 | 0.001 | 0.1 | 0.31 |

**Table S74.** Partial distance correlations and their associated p-values for the different models under analysis, using only non-European traditions, except for "Germans", in the Bronze Age dataset (Supplementary Data, 2.4) and AmAfr motifs (Table SS5).

| Model | Partial correlation | p-value |
|---|---|---|
| Mythemic~genetic, geographic | -0.06 | 0.6 |
| Mythemic~geographic, genetic | 0.47 | 0.004 |

**Table S75.** Model comparison between demic diffusion (mythemic~genetic) and cultural diffusion predicted by isolation-by-distance (IBD) (mythemic~geographic), using only non-European traditions, except for "Germans", in the Iron Age dataset (Supplementary Data, 2.5) and AmAfr motifs (Table SS5). The values reported correspond to Pearson's product-moment correlations (cor), bias-corrected correlations (b.-c. cor), and their respective associated p-values.

| Model | cor | p-value | b.-c. cor | p-value |
|---|---|---|---|---|
| Mythemic~genetic | 0.23 | 0.18 | 0.2 | 0.16 |
| Mythemic~geographic | 0.26 | 0.13 | 0.47 | 0.006 |
| Genetic~geographic | 0.54 | < 0.001 | 0.29 | 0.07 |

**Table S76.** Partial distance correlations and their associated p-values for the different models under analysis, using only non-European traditions, except for "Germans", in the Iron Age dataset (Supplementary Data, 2.5) and AmAfr motifs (Table SS5).

| Model | Partial correlation | p-value |
|---|---|---|
| Mythemic~genetic, geographic | 0.07 | 0.32 |
| Mythemic~geographic, genetic | 0.44 | 0.01 |

**Table S77.** Model comparison between demic diffusion (mythemic~genetic) and cultural diffusion predicted by isolation-by-distance (IBD) (mythemic~geographic), using only non-European traditions, except for "Germans", in the present-day dataset (Supplementary Data, 2.6) and AmAfr motifs (Table SS5). The values reported correspond to Pearson's product-moment correlations (cor), bias-corrected correlations (b.-c. cor), and their respective associated p-values.

| Model | cor | p-value | b.-c. cor | p-value |
|---|---|---|---|---|
| Mythemic~genetic | 0.44 | 0.007 | 0.44 | 0.01 |
| Mythemic~geographic | 0.26 | 0.13 | 0.47 | 0.006 |
| Genetic~geographic | 0.68 | < 0.001 | 0.74 | < 0.001 |

**Table S78.** Partial distance correlations and their associated p-values for the different models under analysis, using only non-European traditions, except for "Germans", in the present-day dataset (Supplementary Data, 2.6) and AmAfr motifs (Table SS5).

| Model | Partial correlation | p-value |
|---|---|---|
| Mythemic~genetic, geographic | 0.16 | 0.19 |
| Mythemic~geographic, genetic | 0.24 | 0.11 |

**Figure S11.** Correlation values over time. We calculated the bias-corrected (b-c. cor) and partial distance correlations (p.cor) between mythological and geographic (cultural model) or genetic (demic model) distances (Materials and Methods) over time for 8 Asian and one European populations, either using all motifs present in the dataset (A) or using only motifs present in at least one African and one South American populations (Table SS5) (B). Unfilled dots indicate non-statistically significant relationships (p-value > 0.05).

A.

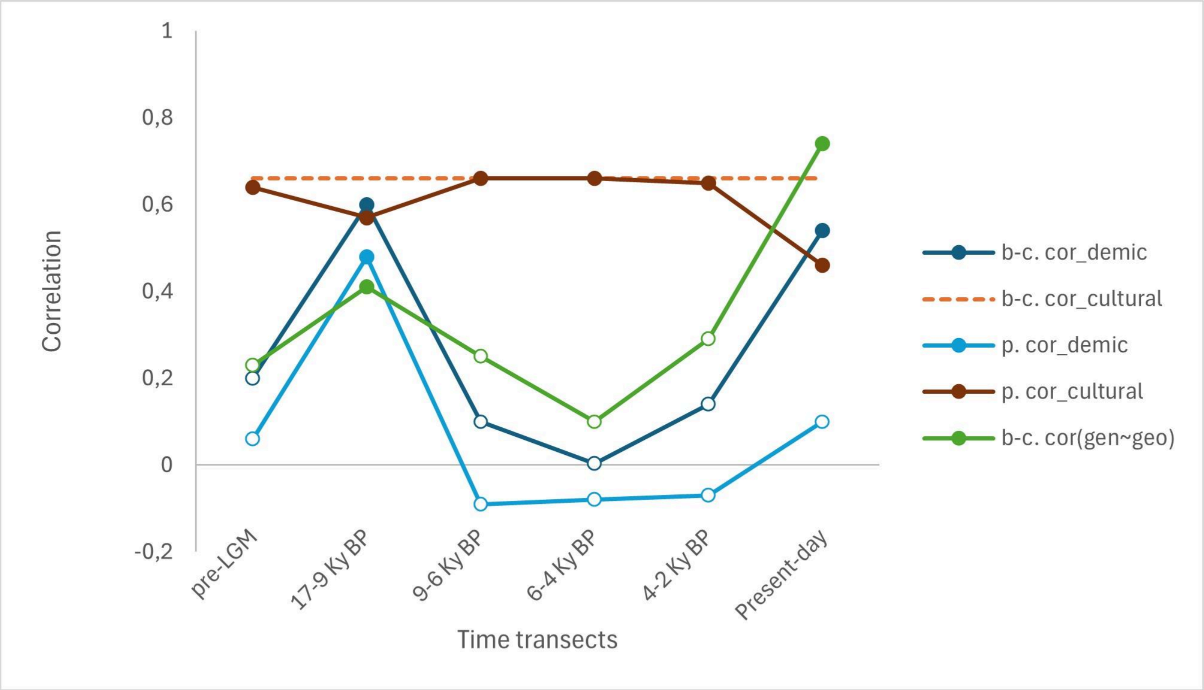


B.

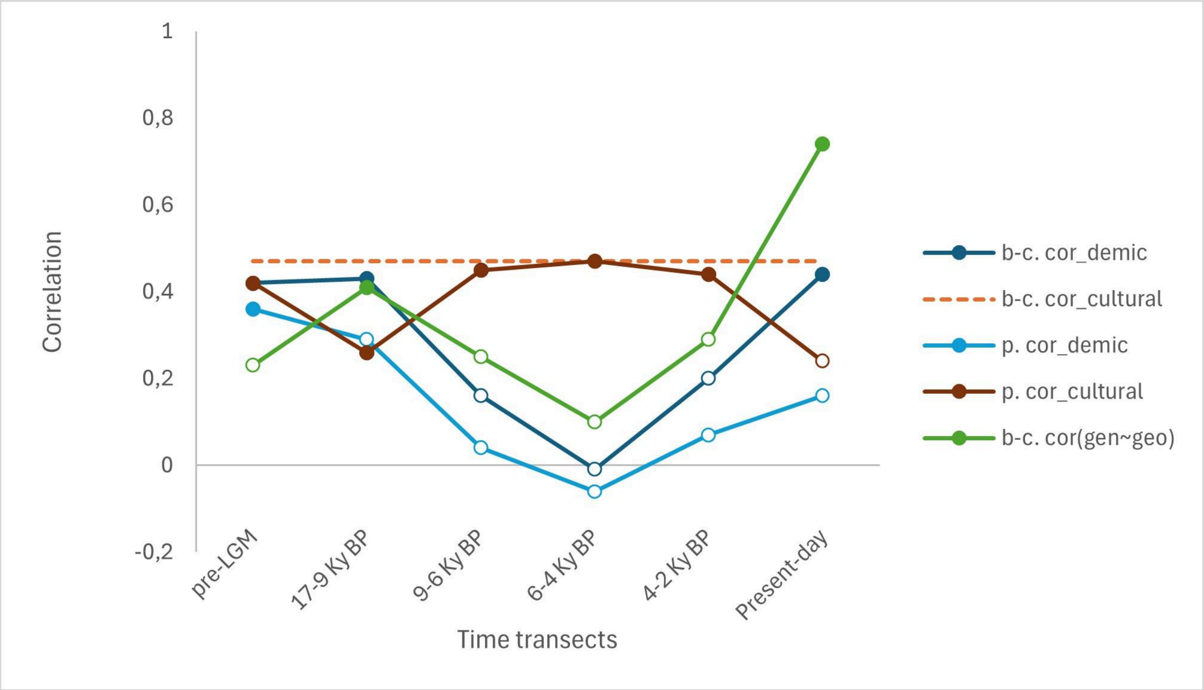

**Identifying particularly demic motifs**

**Table S79.** Motifs associated with a statistically significant linear regression model and genetic regression coefficient, with the latter being positive as well, and with a large genetic effect size measured with omega-squared. Their meanings are taken from Berezkin's database (https://www.mythologydatabase.com/bd/intro.html) consulted on the 11th of January, 2024.

| Motif | Meaning | Description |
|---|---|---|
| b9_3 | Water in the tree trunk. | There is enormous amount of water inside a trunk of a tree or a tree turns into water. |
| g28_6 | The fish tree. | Tree contains fish in its trunk. |
| i41_3 | Rainbow serpent | Rainbow is a reptile (usually a snake) or (more rare) a fish, or it is related to snake, to its tongue, breath, or to scorpion's tail. |
| i82c_2 | Venus is the Moon's wife. | Venus or some other bright star seen near the eastern or western horizon is female and wife of the Moon. |
| j15_10 | Woman gets to dangerous creature. | Walking in search of her husband, boyfriend, kinsmen, shelter woman or girl gets to the house of dangerous creatures where she is injured or killed. |
| L7_10 | Chasing an animal by mistake. | Instead of chasing a person, a bush spirit, a monster or a dangerous animal pursues by mistake an object or animal that moves nearby. |
| m8_10 | Breaking the obstacle. | Non-human persons work hard to destroy a strong and durable obstacle that blocks access to some place or object. |

| | | |
|---|---|---|
| m29v_9 | Trickster is a small ungulate. | In episodes related to deception, absurd, obscene or anti-social behavior the protagonist is a small ungulate (deer, gazelle, antelope). |
| m44b_11 | Thieves of the food: the women. | Person discovers that somebody steals game or fish from his trap or devastates his garden. He or his guards catch the thieves who prove to be (the first) women or the thief is the water being whom the hero lets go after receiving a woman for ransom. |

**Figure S12.** Maps of the worldwide distribution of the motifs associated with a statistically significant (p-value > 0.05) linear model and genetic regression coefficient, the latter being also positive, and associated with a large effect size.

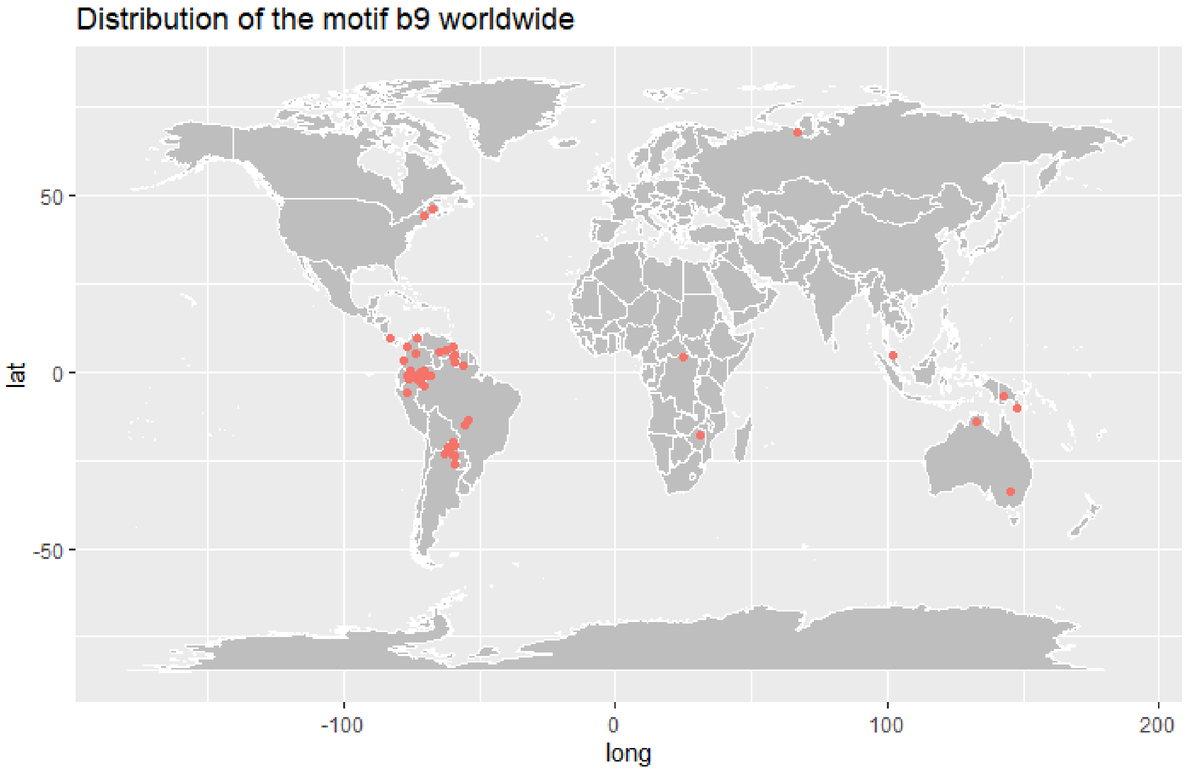

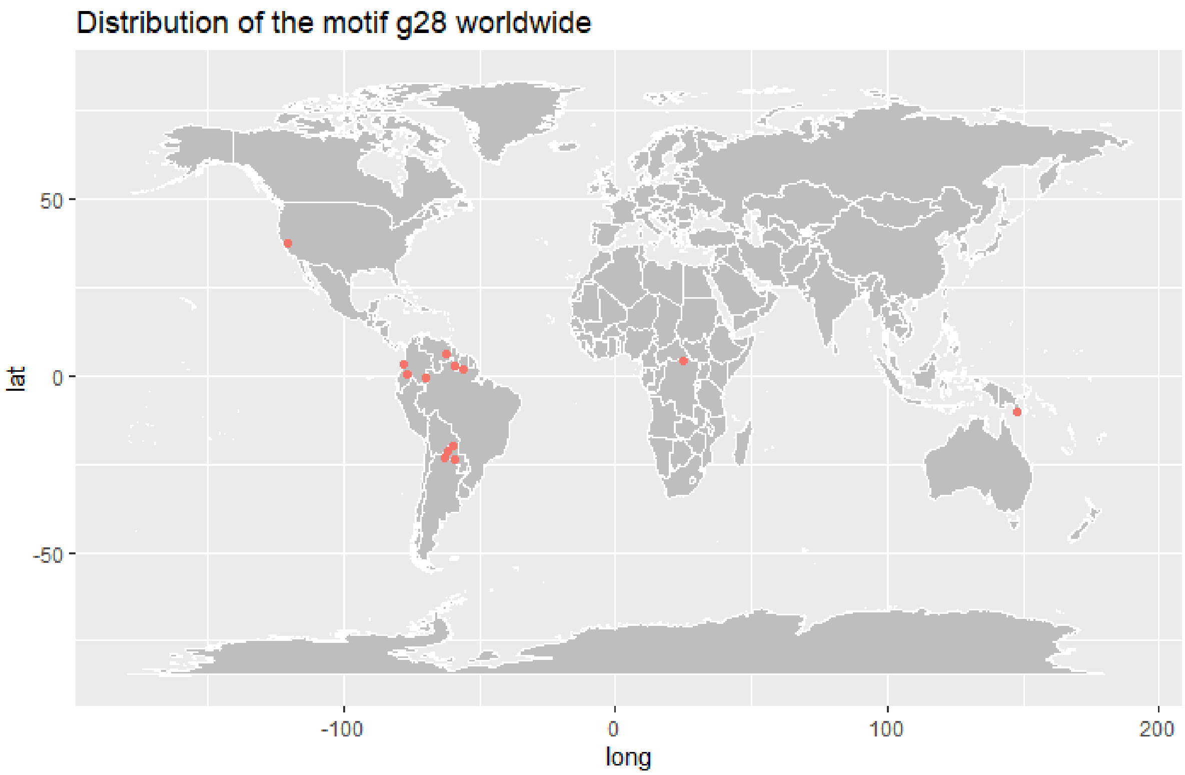
Distribution of the motif g28 worldwide
50
0
-50
lat
-100
0
100
200
long

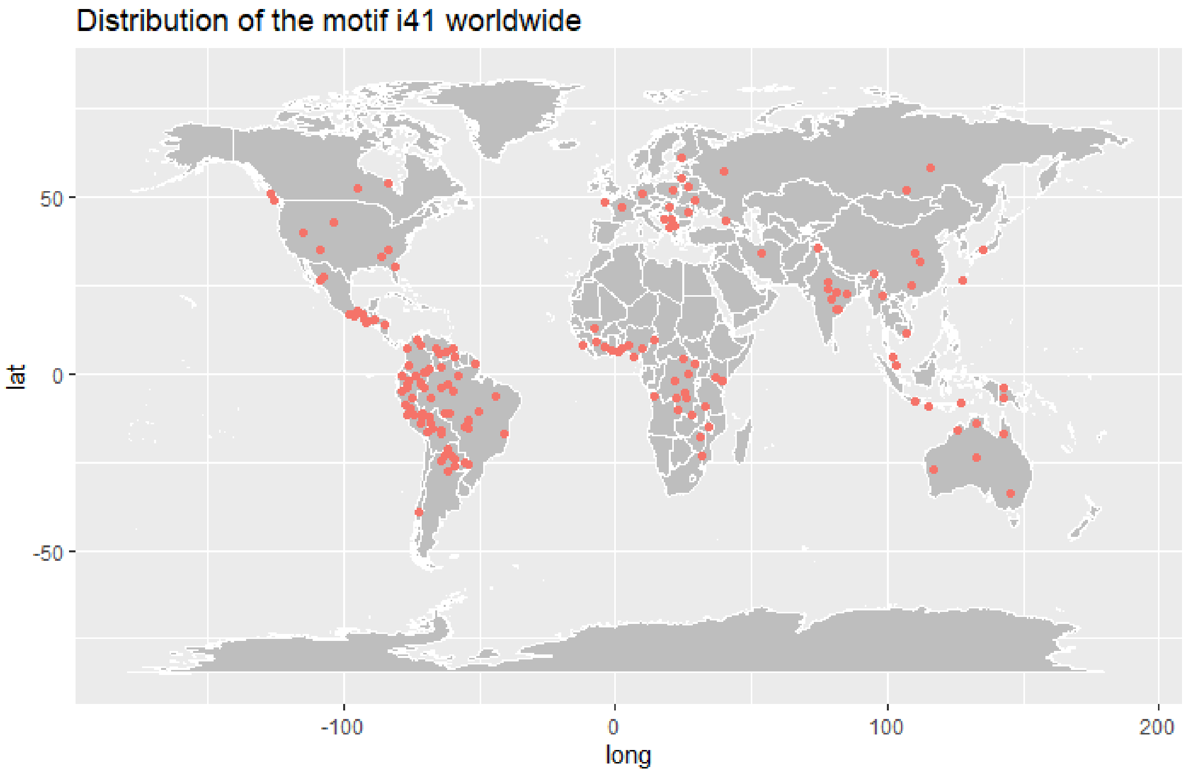
Distribution of the motif i41 worldwide
50
0
-50
lat
-100
0
100
200
long

Distribution of the motif i82c worldwide

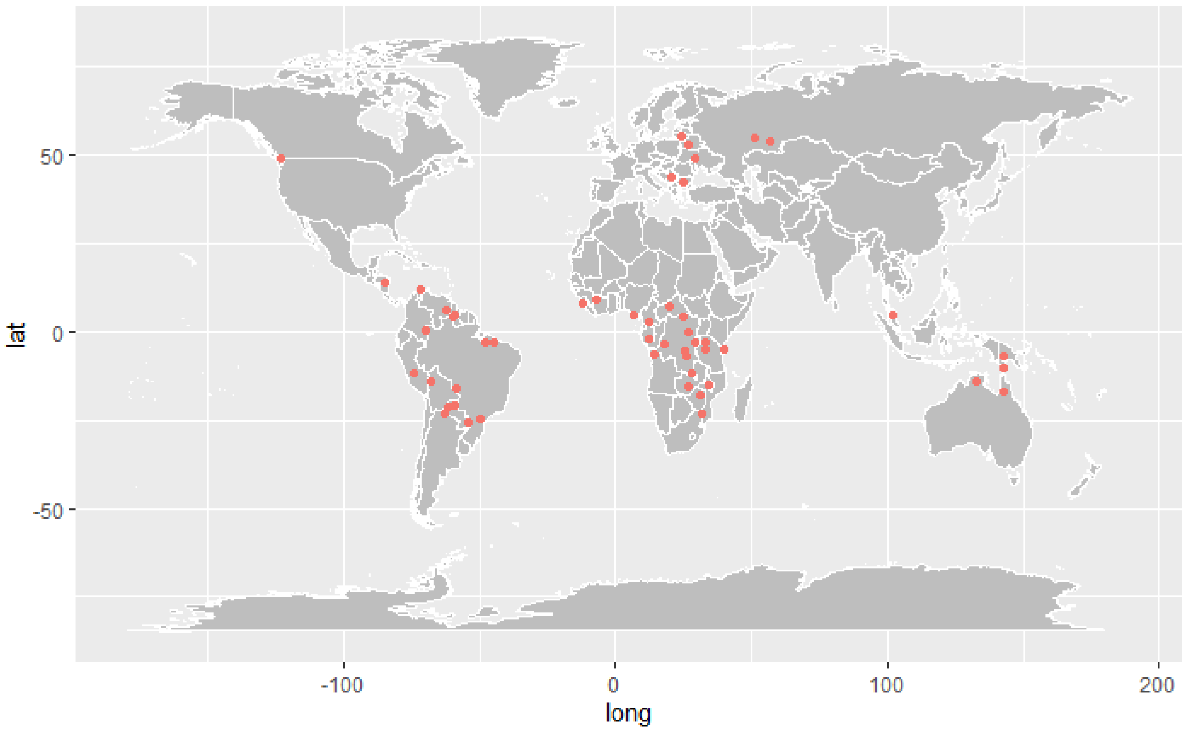


Distribution of the motif j15 worldwide

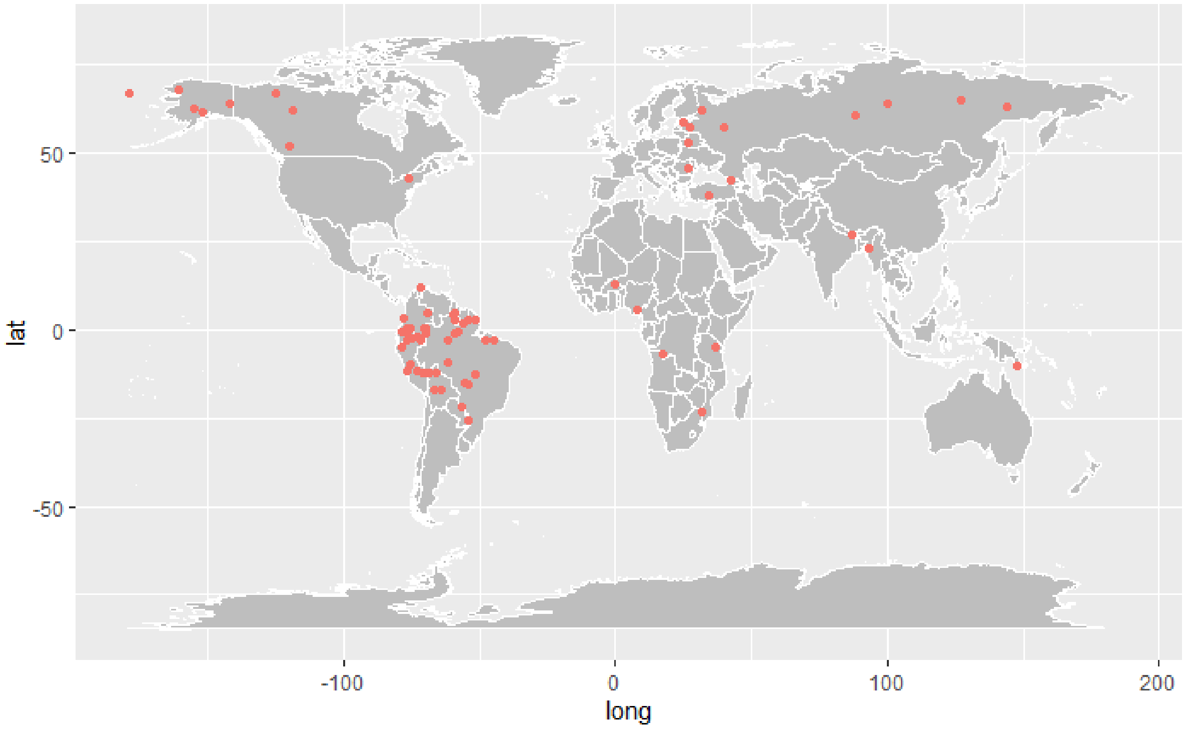

Distribution of the motif L7 worldwide

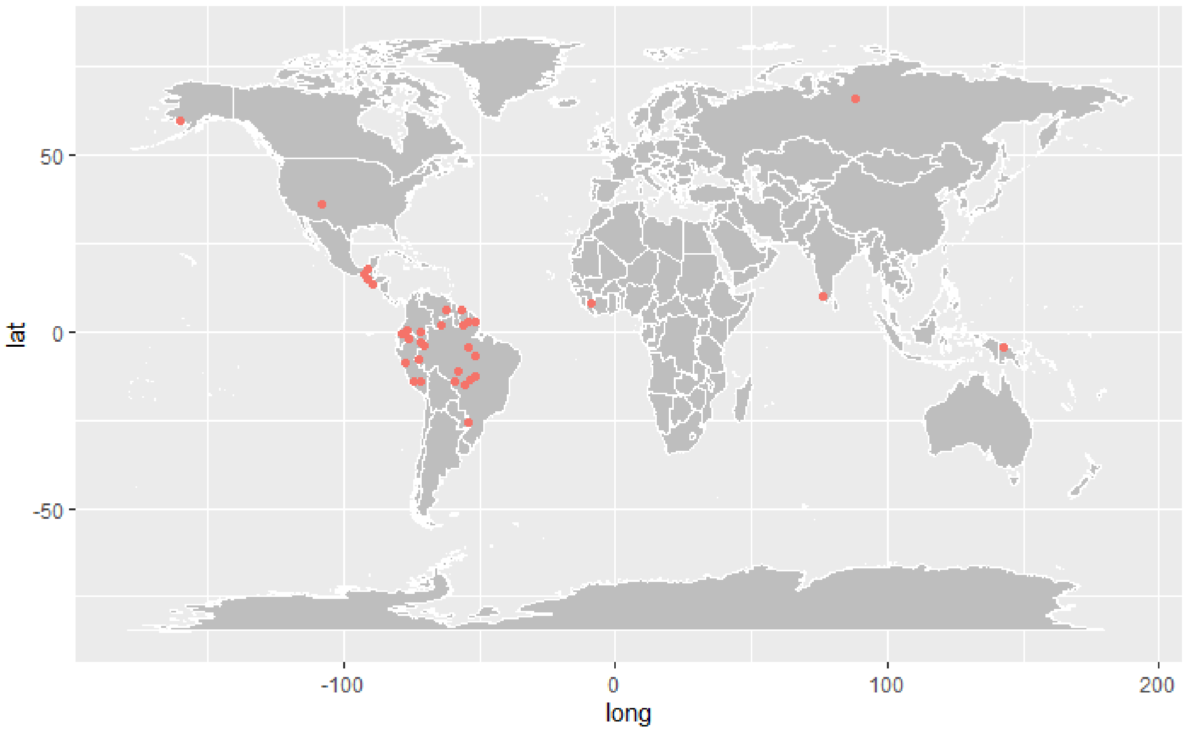


Distribution of the motif m8 worldwide

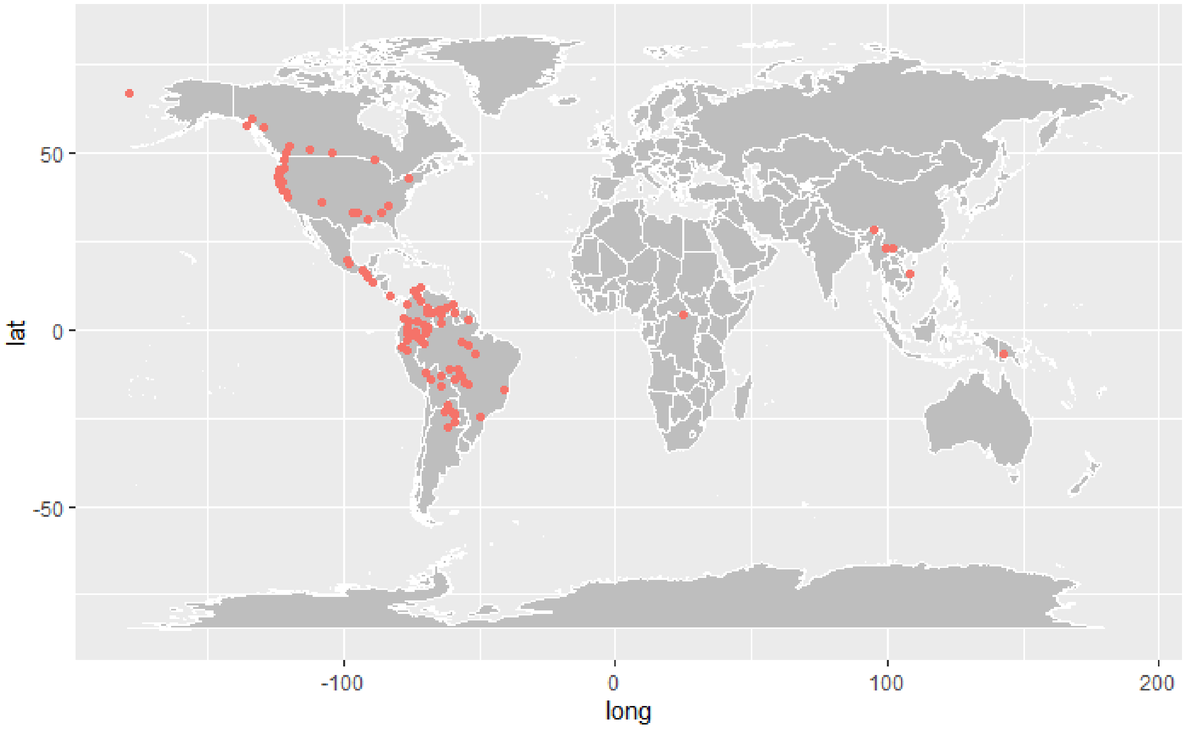

Distribution of the motif m29v worldwide

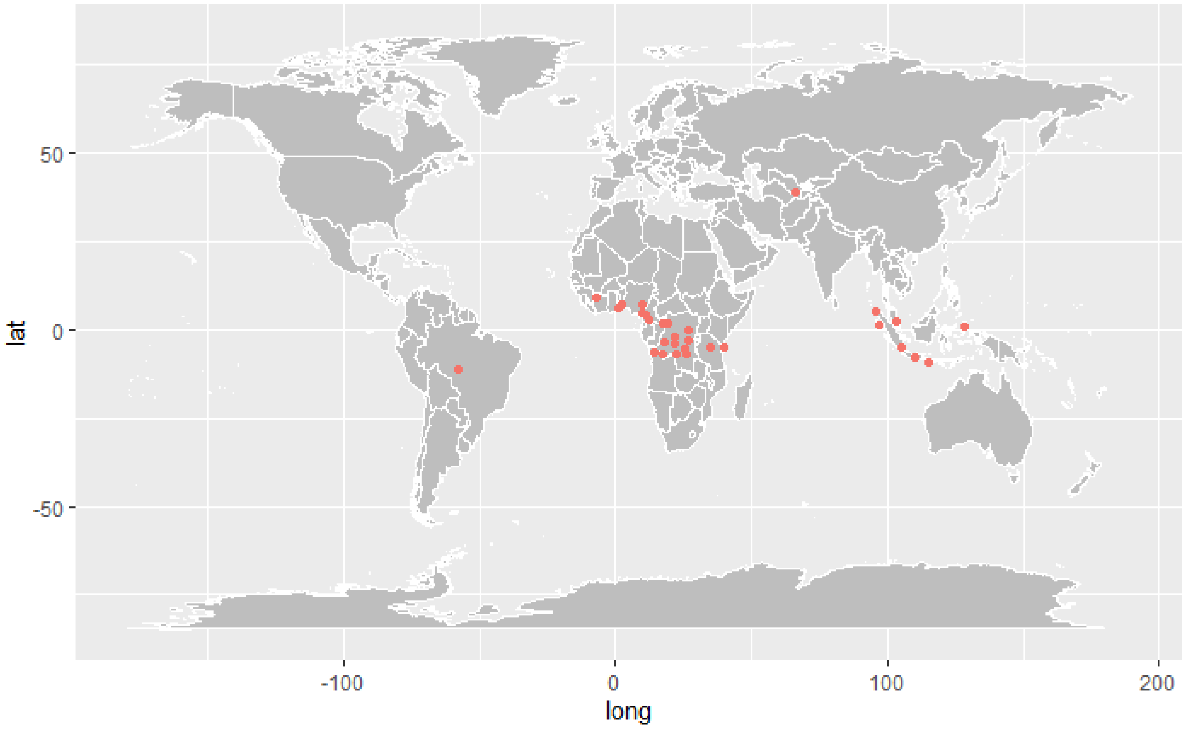


Distribution of the motif m44b worldwide

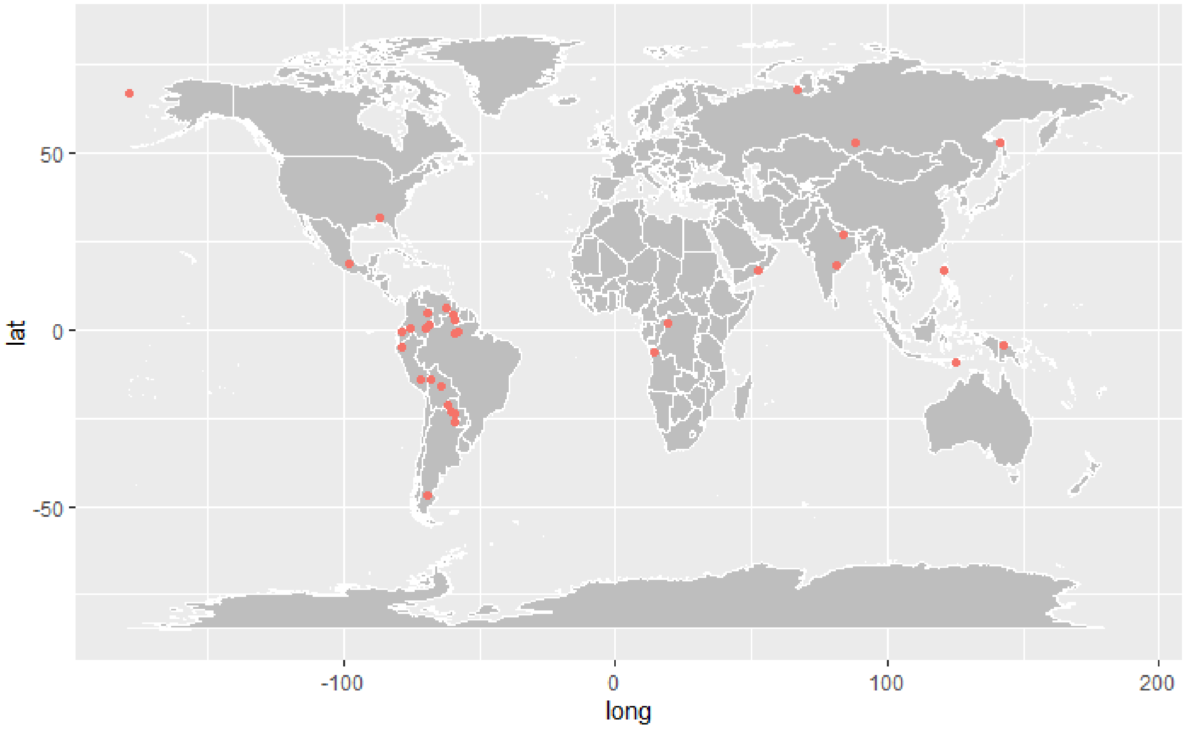

**Biases in mythological database**

Figure S13. Distribution of the number of mythological traditions described in each broad world region.

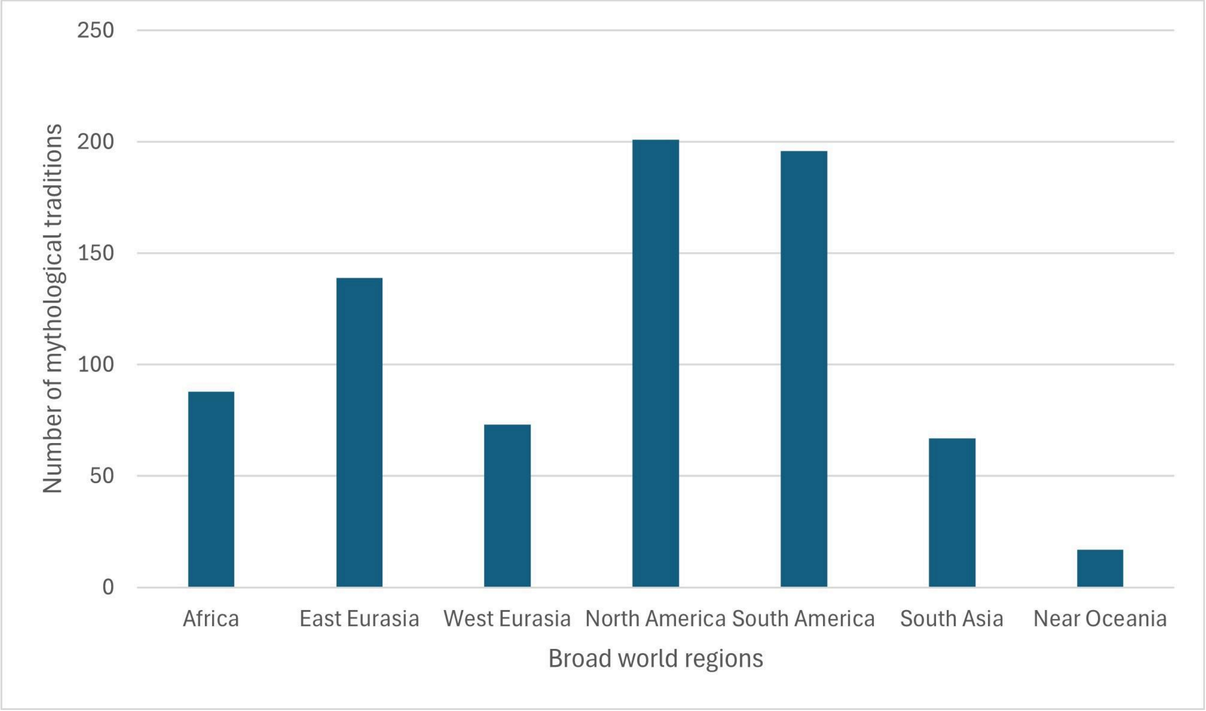


Figure S14. Distribution of the number of motifs described in each tradition present in Berezkin's database, either on a global scale or per world region.

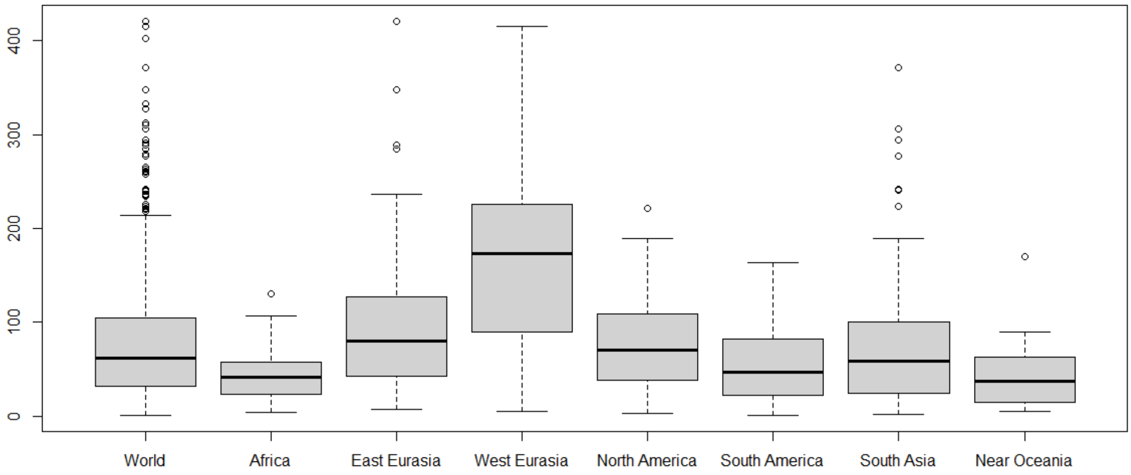

### References


1. Bortolini, E. *et al.* Inferring patterns of folktale diffusion using genomic data. *Proc Natl Acad Sci U S A* **114**, 9140–9145 (2017).
2. d'Huy, J. *Cosmogonies*. (La Découverte, 2020).
3. d'Huy, J. Un récit de plongeon cosmogonique au Paléolithique supérieur ? *Préhistoire du Sud-Ouest* **25**, 109–117 (2017).
4. d'Huy, J. Coda : La symphonie du premier plongeon. *Préhistoire du Sud-Ouest* **26**, 179–190 (2018).
5. d'Huy, J. « Mort, où est ta victoire ? » Reconstruction statistique des premières croyances de l'humanité sur la mort. *Paléo* 182–195 (2020).
6. d'Huy, J. & Berezkin, Y. E. How Did First Humans Perceive Starry Night? On the Pleiades. *Retrospective Methods Network Newsletter* 100–122 (2017).
7. d'Huy, J. Matriarchy prehistory: statistical method testing old theory. *Les Cahiers de l'AARS* **19**, 159–170 (2017).
